# CD28 costimulatory domain protects against tonic signaling-induced functional impairment in CAR-Tregs

**DOI:** 10.1101/2020.11.19.390450

**Authors:** Baptiste Lamarthée, Armance Marchal, Soëli Charbonnier, Tifanie Blein, Emmanuel Martin, Katrin Vogt, Matthias Titeux, Marianne Delville, Hélène Vinçon, Emmanuelle Six, Nicolas Pallet, Dany Anglicheau, Christophe Legendre, Jean-Luc Taupin, Birgit Sawitzki, Sylvain Latour, Marina Cavazzana, Isabelle André, Julien Zuber

## Abstract

The use of chimeric antigen receptor (CAR)-engineered regulatory T cells (Tregs) has emerged as a promising strategy to promote immune tolerance in transplantation. However, in conventional T cells (Tconvs), CAR expression is often associated with tonic signaling resulting from ligand-independent baseline activation. Tonic signaling may cause CAR-T cell dysfunction, especially when the CAR structure incorporates the CD28 costimulatory domain (CSD) rather than the 4-1BB CSD.

Here, we explored the impact of tonic signaling on human CAR-Tregs according to the type of CSD. Compared to CD28-CAR-Tregs, 4-1BB-CAR-Tregs showed enhanced proliferation and greater activation of MAP kinase and mTOR pathways but exhibited decreased lineage stability and reduced abilities to produce IL-10 and be restimulated through the CAR. Although both CAR-Treg populations were suppressive *in vivo,* cell tracking with bioluminescence imaging found longer persistence for CD28-CAR-Tregs than for 4-1BB-CAR-Tregs. This study demonstrates that CD28-CAR best preserves Treg function and survival in the context of tonic signaling, in contrast with previous findings for Tconvs.

**SUMMARY:** Lamarthée et al investigated the impact of chimeric antigen receptor (CAR) tonic signaling on CAR-engineered Tregs according to the incorporated costimulatory domain (either CD28 or 4-1BB). CD28 ameliorated Treg stability, long-term survival and function compared to 4-1BB.

## INTRODUCTION

Despite significant advances in the prevention of acute graft rejection, long-term attrition of graft function and complications related to long-term immunosuppression remain major concerns in the field of solid organ transplantation. In addition to their role in the control of self-reactivity (Zuber et al., 2007), FOXP3-expressing regulatory T cells (Tregs) play a key role in mitigating alloimmune responses in experimental (Kendal et al., 2011) and clinical transplantation (Savage et al., 2018). Hence, significant efforts have been made to apply regulatory T cell therapy in clinical transplantation over the past decade (Sawitzki et al., 2020). However, although experimental transplant models indicate that donor-specific Tregs produce greater prevention of graft rejection than polyclonal Tregs (Sagoo et al., 2011), the generation of clinical-grade donor-specific Tregs faces major challenges.

These include the isolation of the scarce antigen-specific population and its subsequent expansion to produce a sizeable cell product (Sagoo et al., 2011). An attractive alternative would be to engineer recipient Tregs with a chimeric antigen receptor (CAR) that redirects their antigen specificity toward a given donor antigen. CARs are engineered receptors that consist of an extracellular single-chain variable fragment (scFv), which is derived from the antigen-binding regions of a monoclonal antibody, joined to intracellular T cell signaling domains called cluster of differentiation (CD)3ζ combined with a costimulatory domain (CSD), most frequently that of either 4- 1BB or CD28.

A few pioneering studies have provided proof-of-concept evidence that the Treg response can be redirected toward a donor mismatched antigen. More specifically, Human Leukocyte Antigen (HLA) A2-targeted CAR-Tregs efficiently prevent graft-versus-host disease (GVHD) and skin graft rejection in an HLA-A2-specific manner (Boardman et al., 2017; Boroughs et al., 2019; MacDonald et al., 2016; Noyan et al., 2017; Pierini et al., 2017; Dawson et al., 2020). Most of these pilot studies have used HLA-A2-targeted CARs incorporating the CD28 endodomain based on evidence that the CD28 signal is critical for Treg ontogeny (Salomon et al., 2000; Watanabe et al., 2005), survival (Zhang et al., 2013) and epigenetic stability (DuPage et al., 2015). On the other hand, 4-1BB expression is a hallmark marker of activated human Tregs (Bacher et al., 2016; Schoenbrunn et al., 2012), and *TNFRSF9* (Tumor Necrosis Factor Receptor SuperFamily 9, encoding 4-1BB) belongs to a handful of genes whose epigenetic marks are highly conserved in activated Tregs across species (Arvey et al., 2015). Emerging data suggest that the 4-1BB CSD could support greater proliferation than its CD28 counterparts at the expense of Treg suppressive function (Boroughs et al., 2019; Dawson et al., 2020; Nowak et al., 2018).

In the field of oncology, studies have shown that tonic activation of CARs, i.e., signaling irrespective of the presence of the CAR ligand, occurs to varying degrees with most CARs, resulting in baseline activation and eventually leading to T cell exhaustion with a reduced antitumor effect (Long et al., 2015). Importantly, the 4-1BB CSD was found to significantly reduce tonic signaling-induced exhaustion and be associated with longer survival than the CD28 CSD (Long et al., 2015). In the field of CAR-Tregs, however, it is largely unknown whether CAR-mediated tonic signaling impacts Treg biology and fate and whether the type of CSD (4-1BB vs CD28) can influence this effect.

In the present study, we exploited the tonic signaling induced by an anti-HLA-A2 CAR to assess its effect on CAR-Treg phenotype, proliferation, metabolism, signaling and function, according to the type of CSD. 4-1BB CAR tonic signaling resulted in enhanced expansion following TCR stimulation but was associated with Treg destabilization upon long-term culture. A Treg-supporting cocktail, including a mechanistic target of rapamycin (mTOR) inhibitor, could curb Treg proliferation and promote FOXP3 expression. However, 4-1BB CAR-Tregs still exhibited high levels of baseline activation (HLA-DR expression), a relatively short *in vivo* lifespan and a reduced ability to be stimulated through the CAR. We concluded that in contrast to previous findings for CAR-T cells (Long et al., 2015), the CD28 CSD was more suitable than the 4-1BB CSD for CAR-Treg engineering, especially in the context of tonic CAR signaling.

## RESULTS

### Generation of CD28 and 4-1BB HLA-A2-specific CAR-Tregs

Six anti-HLA-A2 scFvs were derived from the publicly available sequences of three human anti-HLA-A2/A28 antibodies (Watkins et al., 2000). To assess their specificity/affinity, the 6 scFvs were tagged with a polyhistidine tail and then mixed with 96 beads, each of which was coated with a single HLA class I antigen (Figure Supp 1A). One scFv was selected based on its exclusive binding to beads coated with HLA-A2 and HLA-A28 antigens (Figure Supp 1B). To assess the impacts of the selected CSD on CAR-Treg biology and function, two anti-HLA-A2 CAR constructs were generated with the selected scFv, incorporating either the CD28 CSD or the 4-1BB CSD, named thereafter CD28 CAR and 4-1BB CAR, respectively (Figure 1A). A reporter gene encoding truncated epidermal growth factor receptor (EGFRt) was placed behind thosea asigna virus 2A (T2A). Both bicistronic constructs were next incorporated into a pCCL lentiviral vector (LV) backbone. To demonstrate that antigen specificity was maintained after scFv incorporation into the CAR structure, TCR-deficient J.RT3-T3.5 Jurkat cells were transduced with one of the HLA-A2-specific CARs and then challenged against a panel of irradiated splenocytes isolated from 10 HLA-typed deceased organ donors. Induction of CD69 expression in CAR-expressing Jurkat cells was strictly restricted to those cocultured with irradiated splenocytes expressing either HLA-A2 antigens or HLA-A28 antigens (Figure Supp 1C).

**Figure 1:**
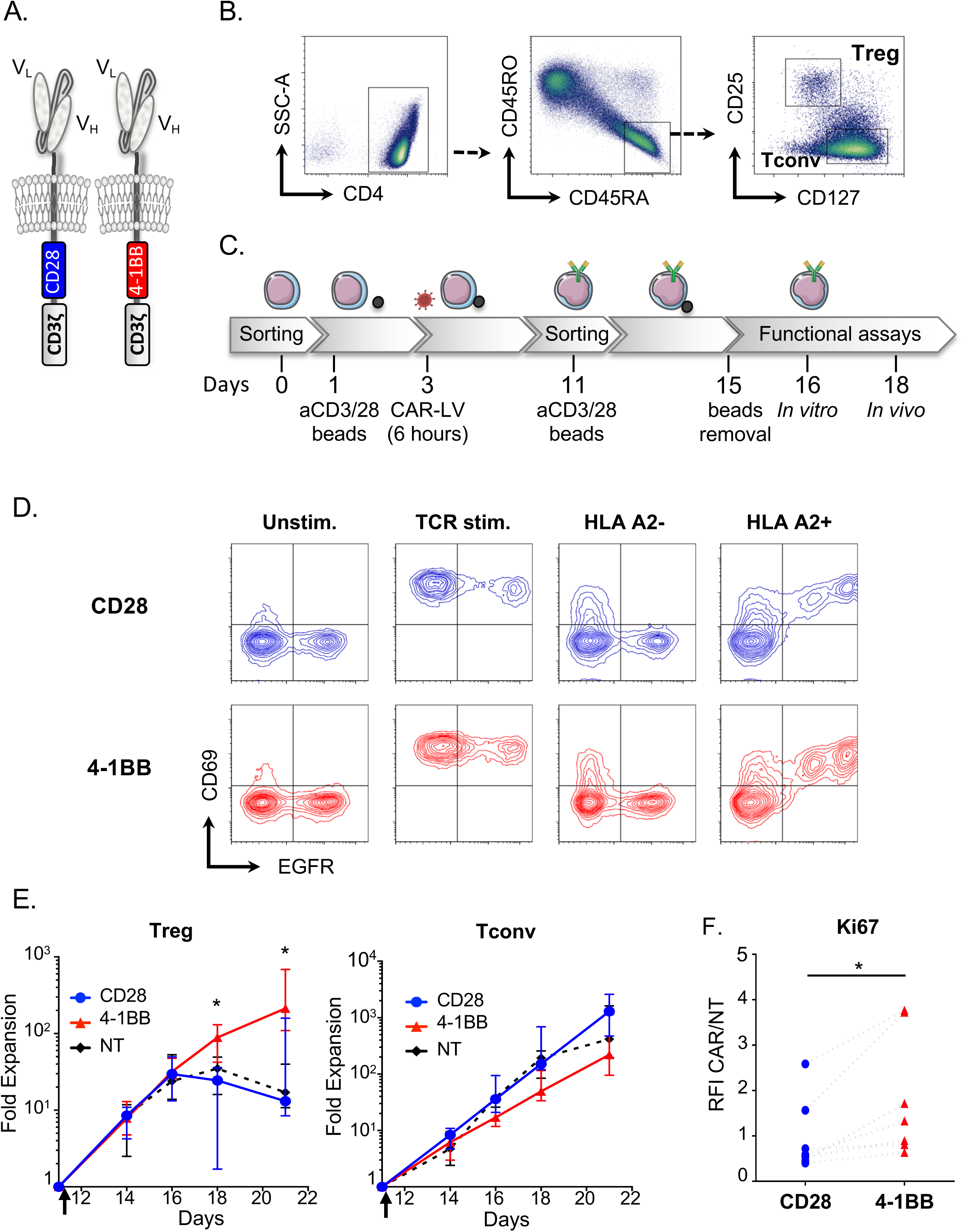
CD28 and 4-1BB CAR-Treg production. A. Schematic representation of CD28 and 4-1BB CARs. B. Gating strategies for CD4+ CD45RA+ CD45RO-CD25hi CD127low naïve Tregs and CD4+ CD45RA+ CD45RO-CD25low CD127hi Tconvs sorted by flow cytometry from enriched HLA-A2-CD4+ PBMCs. CD4+ Tregs and Tconvs were selected on morphology and exclusion of doublets and dead cells (not shown) before positive selection. C. Schematic depicting the process for CD28 or 4-1BB CAR-T cell production. T cells were isolated, stimulated, transduced with CAR-encoding lentiviruses, expanded, sorted based on reporter gene expression, restimulated and expanded for up to 21 days. D. On day 11, CD28 and 4-1BB CAR-Tregs were incubated with or without CD3/CD28 beads or HLA-A2-positive or HLA-A2-negative irradiated splenocytes, and CD69 and EGFR expression was measured by FACS. Contour plots are representative of 2 experiments. E. Fold expansion of CD4+ Tregs and untransduced Tregs (left panel) or their respective Tconv populations (right panel) from TCR restimulation on day 11 to day 21 of culture. The median+/- interquartile range (IQR) is represented. N=10-15 donors, pooled from at least 10 independent experiments. Mann-Whitney test at the indicated time points comparing 4-1BB and CD28 CAR-Tregs, *p<0.05. F. Ki67 expression measured by flow cytometry on day 16 of culture. The ratio of fluorescence intensity (RFI) was calculated as follows: the geometric mean of fluorescence (MFI) of either CD28 or 4-1BB CAR-Tregs divided by the MFI of untransduced Tregs. Each replicate is represented. N=7 donors from 7 independent experiments. Wilcoxon matched-pairs signed rank test *p<0.05.

Since naïve Tregs have been found to be more stable over long-term culture than activated Tregs (Hoffmann et al., 2006; Miyara et al., 2009), CD4+ CD25^Hi^ CD127-CD45RA+ CD45RO- naïve Tregs and CD4+ CD25-CD127+ CD45RA+ CD45RO-naïve conventional T cells (Tconvs), which served as controls, were sorted by fluorescence-activated cell sorting (FACS) with HLA-A2/A28-negative cytapheresis kits from young (<40 years old) healthy donors (Figure 1B). Tregs and Tconvs were transduced with HLA-A2-targeted CAR-encoding LV vectors after 2 days of polyclonal activation with CD3/CD28 beads. EGFRt-expressing transduced T cells were sorted by FACS and restimulated on day 11 before further *in vitro* (day 16) and *in vivo* (day 18) assessment (Figure 1C). To further evaluate HLA-A2-specific activation of CAR-engineered primary T cells, CAR-Tregs were collected on day 10 and, 72 hours after bead separation, were restimulated with HLA-A2/A28-positive or HLA-A28-negative irradiated splenocytes. CD69 (Figure 1D) and Glycoprotein-A Repetitions Predominant (GARP) (not shown) expression was strongly induced in an HLA-A2-specific manner by EGFR-expressing Tregs but not by untransduced EGFR-negative Tregs. These results indicate that both vectors were efficiently expressed on the cell surface of human Tregs and that CAR stimulation was able to induce antigen-specific activation of CAR-expressing human Tregs.

### 4-1BB induces strong tonic signaling, which enhances Treg proliferation

In line with a recent report (Dawson et al., 2020), we observed that the proliferative capacity of CAR-Tregs greatly varied between CD28 CAR-Tregs and 4-1BB CAR-Tregs. Indeed, 4-1BB CAR-Tregs had achieved 20-fold greater expansion than their CD28 counterparts at day 21. Furthermore, 4-1BB-Treg expansion did not plateau even after 3 weeks of culture, whereas other Treg populations stopped proliferating shortly after the second CD3/CD28 stimulation (Figure 1E, left panel). These results were Treg specific, as 4-1BB CAR-Tconvs and CD28 CAR-Tconvs showed similar growth profiles (Figure 1E, right panel). Consistent with this heightened and sustained proliferation, 4-1BB CAR-Tregs demonstrated significantly higher expression of Ki67 than CD28 CAR-Tregs (Figure 1F).

Notably, unlike CD28 CAR-Tregs, 4-1BB CAR-Tregs showed a blastic phenotype throughout cell culture, as observed by optical microscopy (not shown) and flow cytometry side and forward scatter patterns (Figure 2A-B). However, in contrast to previous studies, we found that the 4-1BB CSD had a positive effect on Treg proliferation when CAR-Tregs were stimulated through their endogenous TCR. This indicated that 4-1BB-driven enhanced proliferation was constitutive and ligand independent and could therefore be ascribed to tonic signaling (Ajina and Maher, 2018; Frigault et al., 2015; Gomes-Silva et al., 2017). Ligand-independent CAR tonic signaling seems to primarily depend on the high cell-surface density and intrinsic characteristics (scFv and hinge/spacer domains) of the CAR, which altogether entail CAR clustering (Ajina and Maher, 2018). Hence, we compared CAR expression at the cell surface according to the CSD and cell lineage. Although the vector copy number (VCN) in transduced Tregs was similar between CD28 CAR and 4-1BB CAR (Figure 2C), 4-1BB CAR-Tregs displayed stronger binding to the HLA-A2 pentamer and greater EGFR expression than CD28 CAR-Tregs and CAR-Tconvs (Figure 2B). This flow cytometry pattern might indicate greater vector expression and higher CAR and EGFR protein production at the cellular level in 4-1BB CAR-Tregs. However, this pattern was tightly correlated with the blastic phenotype (Figure 2B). When normalized to the cell surface area (as estimated by the squared forward scatter pattern, FSC^2^), the mean fluorescence intensities (MFIs) of HLA-A2 pentamer and EGFR staining were not different across the CAR CSDs and cell lineages (Figure 2D). We next sought to measure CAR expression, independent of the ability of each CAR to bind HLA-A2, as this could be a confounding variable across the different CARs. To this end, CAR-Tregs were labeled with biotinylated protein L, a recombinant protein that binds to the kappa light chain without interfering with the antigen-binding site. Similarly, normalized CAR expression, as assessed by protein L staining, was not significantly different across the CAR CSDs and T cell subsets (Figure 2D, right panel). We inferred that 4-1BB tonic signaling, primarily observed in Tregs, could not be ascribed to a greater transduction efficacy or to an increased CAR cell-surface density. Instead, these results suggest that constitutive 4-1BB stimulation itself interferes with human Treg biology.

**Figure 2:**
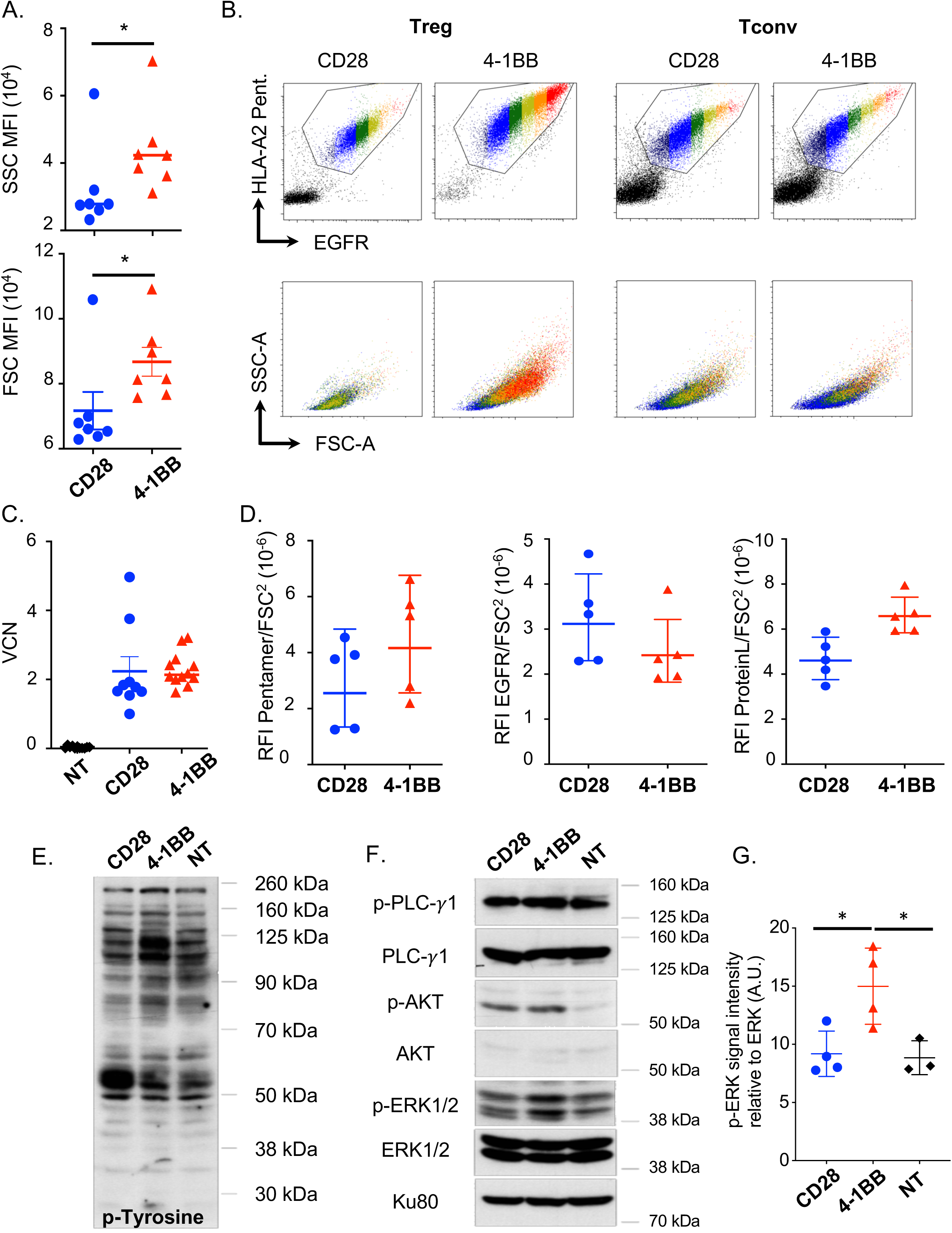
4-1BB tonic signaling induces strong CAR-Treg proliferation associated with MTOR and MAPK pathway activation. A. Flow cytometry side (upper panel) and forward (lower panel) scatter parameter MFIs of 4-1BB and CD28 CAR-Tregs on day 11 of culture. N=7 donors from 7 independent experiments. Mann-Whitney test, *p<0.05. B. Flow cytometry analysis of HLA-A2 pentamer binding and EGFRt reporter gene expression (upper panels) and FSC-SSC parameters (lower panels). A color scale was applied to gate transduced cells. Dot plots are representative of 7 experiments. C. Vector copy number on day 16 of culture. Each replicate and the medians are represented. N=9 donors from 9 independent experiments. D. Ratios of pentamer MFI (left panel), EGFR MFI (middle panel) and protein L MFI (right panel) normalized by CAR-Treg cell surface are (estimated by as the squared FSC). Each replicate and the medians of 4 independent experiments are represented. E. Immunoblots with antibodies against tyrosine-phosphorylated residues. F. Phosphorylated PLC-γ1, phosphorylated ERK1/2, phosphorylated Akt, and Ku80, a loading control, are shown. Molecular weights are shown on the right. The immunoblot is representative of 4 independent experiments. G. The phospho-ERK signal intensity relative to the ERK signal intensity is depicted in arbitrary units (A.U.) for 4 independent experiments.

### 4-1BB tonic signaling increases MAPK and mTOR pathway activation

We thus sought to further investigate the underlying mechanisms and consequences of CAR tonic signaling in Tregs. During the culture process, both CD28 CAR-Tregs and 4-1BB CAR-Tregs were restimulated with CD3/CD28 beads on day 11 of culture (Figure 1C). On day 16, the 4-1BB CAR-Tregs were still expanding, whereas the CD28 CAR-Tregs had started to plateau (Figure 1E). We thus performed an analysis of tyrosine phosphorylation signals by western blot analysis at this time to better characterize 4-1BB and CD28 tonic signals five days after TCR restimulation. Strikingly, we observed that signaling pathways were markedly different among 4-1BB CAR-Tregs, CD28 CAR-Tregs and untransduced Tregs (Figure 2E). The tyrosine phosphorylation of several proteins was found to be increased, including proteins with molecular weights of 115-120 kDa and 55 kDa in the cell lysates of 4-1BB CAR-Tregs and CD28 CAR-Tregs, respectively, when compared with those of untransduced Tregs. Of note, the total protein amounts were comparable, as revealed by Ku80 expression levels (Figure 2F). Further analyses of specific pathways revealed that the phosphorylation of ERK 1/2 was greater in 4-1BB CAR-Tregs than in CD28 CAR-Tregs and untransduced Tregs, indicating overactivation of the downstream MAP kinase pathway (Figure 2F-G). Moreover, Akt phosphorylation on Ser473 was also increased in CAR-Tregs compared to untransduced cells irrespective of the CSD (Figure 2F). Altogether, these results show that CAR tonic signaling differentially impacts Treg biology depending on the CSD incorporated in the CAR. They also suggest that the sustained proliferation of 4-1BB CAR-Tregs could result from tonic activation of the MAP kinase pathway.

### 4-1BB and CD28 tonic signaling induced different Treg differentiation pathways

We next explored whether the differential effects of CD28 and 4-1BB tonic signaling on CAR-Treg proliferation would have similar impacts on the CAR-Treg phenotype. To compare the effects of the 4-1BB and CD28 CSDs on CAR-Tregs, we used 2-dimensional gating as follows: morphology, single cells, and FVD-live cells. We then applied the t-distributed stochastic neighbor embedding algorithm (tSNE) to visualize high-dimensional cytometry data on a 2-dimensional map at single-cell resolution (Amir et al., 2013). Figure 3A shows the tSNE map for a pool of CD28 CAR-Treg and 4-1BB CAR-Tregs from three different donors. Strikingly, the CD28 CAR-Treg and 4-1BB CAR-Treg populations were highly clustered and separated from one another in a dichotomous way, suggesting a major effect of the CSD on the Treg phenotype. Of note, HLA-DR was mainly expressed by the 4-1BB CAR-Tregs, whereas CD15s expression was limited to a small subset of HELIOS-negative CD28 CAR-Tregs. The CD28 CAR-Tregs maintained CCR7 expression overall, in contrast to their 4-1BB counterparts. Interestingly, a sizeable population of the 4-1BB CAR-Tregs expressed high levels of FOXP3 and HELIOS along with markers of activation (HLA-DR, 4-1BB, and ICOS) and suppressive functions (TIGIT and CTLA-4). This subset, which was absent in the CD28 CAR-Tregs, met the phenotypic criteria of effector Tregs.

**Figure 3:**
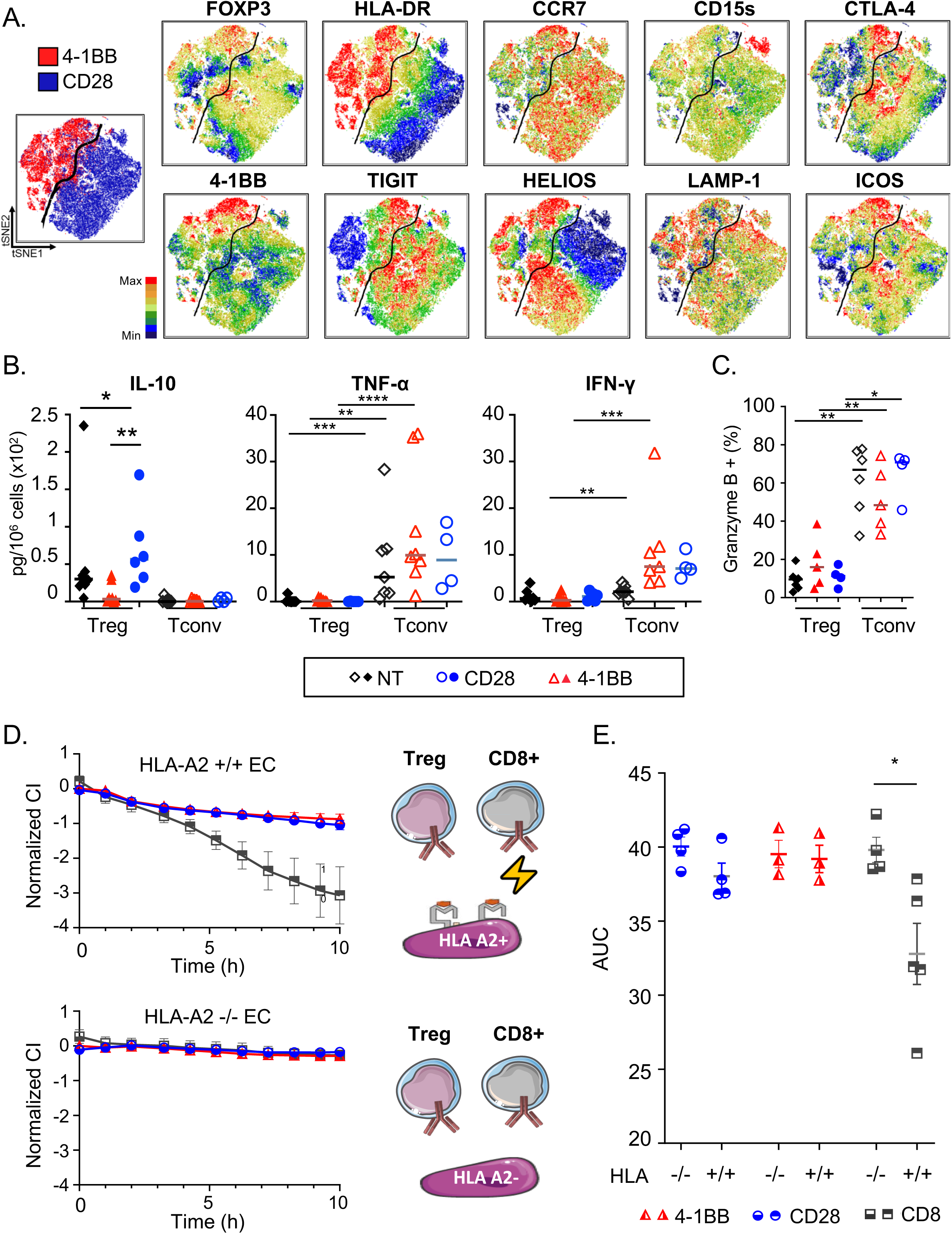
CD28 and 4-1BB CSDs induce different Treg phenotypes. A. tSNE maps of CD28 and 4-1BB CAR-Tregs on day 16 are shown. Each point in the tSNE maps represents an individual cell, and cells are colored according to the intensity of expression of the indicated markers. n=3 donors pooled from three independent experiments. B. On day 14, the amounts of the indicated cytokines were determined by cytometric bead array analysis of culture supernatants. The results were normalized by the number of cells in the corresponding wells at the time of harvest. n=4-9 donors from at least four independent experiments. C. On day 16, cells were TCR-stimulated overnight. The percentage of granzyme B-positive live cells was determined by flow cytometry. D-E. HLA-A2+ endothelial cells (ECs) or HLA class I-deficient ECs were cocultured with CD28 or 4-1BB CAR-Tregs or cytotoxic CD8+ Tconvs as a control. D. The normalized cell index (mean ± standard error) and E. areas under the curve (AUCs) from four independent experiments are shown. Mann-Whitney tests. *p<0.05, **p<0.01, ***p<0.001, ****p<0.0001

### Neither 4-1BB CAR-Tregs nor CD28 CAR-Tregs converted into inflammatory and/or cytotoxic cells

A recent study raised the concern that 4-1BB CARs could switch on a cytotoxic program in human Tregs (Boroughs et al., 2019). Although cytotoxic pathways, including the Fas/FasL and granzyme A/B pathways, are part of Treg-mediated suppressive mechanisms (Vignali et al., 2008), Tregs are not considered predominantly cytotoxic, as their cytotoxic activity is much lower than that of their conventional T cell counterparts. We thus addressed this issue by studying the cytokine production and cytotoxicity of CAR-Tregs on day 14 of culture (Figure 3B) according to the CSD. First, none of the CAR-Treg populations produced inflammatory cytokines (TNF-*α* or IFN-*γ*), in contrast to CAR-Tconvs. It is, however, worth noting that 4-1BB CAR-Tregs produced significantly less IL-10 than CD28 CAR-Tregs (Figure 3B). Second and foremost, granzyme B expression was significantly lower in CAR-Tregs than in CAR-Tconvs, irrespective of the CSD (Figure 3C). To further explore the cytotoxic potential of CAR-Tregs, we set up an *in vitro* impedance-based cytotoxicity assay in which an HLA-A2+ endothelial cell line was used as the target and compared with an HLA class I-deficient counterpart. Importantly, CAR-Tregs did not induce HLA-A2-specific cytotoxicity over 10 hours, regardless of the CSD, in contrast to CD8+ CAR-Tconvs (Figure 3D-E). Overall, these results did not support a drift in CAR-Tregs, regardless of the CSD, toward a cytotoxic/proinflammatory program.

### 4-1BB-driven mTOR pathway overactivation may destabilize the Treg phenotype

We reasoned that increased proliferation of 4-1BB CAR-Tregs would be associated with immunometabolic reprogramming, which would be needed to support the increased anabolic demands. We thus studied the expression of a few genes involved in main metabolic pathways, such as glycolysis, glutaminolysis and purine/pyrimidine synthesis, as previously reported (Fernández-Ramos et al., 2017). We consistently found greater expression of Transition Protein 1 (TP1) and solute carrier family 1 (neutral amino acid transporter) member 5 (SLC1A5) transcripts in 4-1BB CAR-Tregs than in CD28 CAR-Tregs, suggestive of activated glycolytic and glutaminolytic pathways, respectively (Figure 4A). These findings were further confirmed by FACS analysis, which showed augmented expression of the glutamine transporter ASCT2 encoded by SLC1A5 in 4-1BB CAR-Tregs compared to CD28 CAR-Tregs (Figure 4B). The higher glutamine intake by 4-1BB CAR-Tregs was in keeping with the increased proliferation rate.

**Figure 4:**
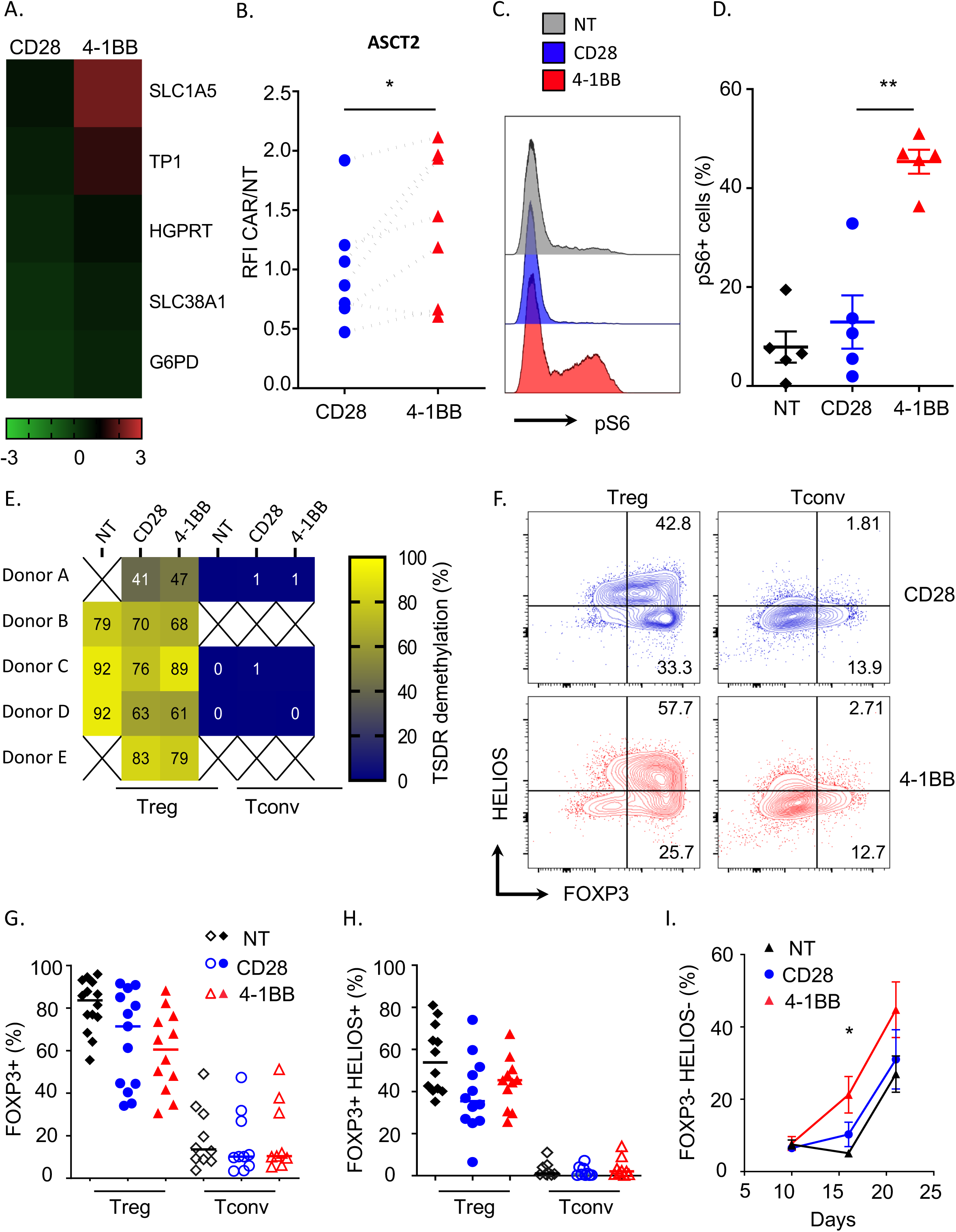
4-1BB CSD drives mTOR pathway overactivation and destabilizes the Treg phenotype upon long-term culture *in vitro*. A. On day 16, Tregs were harvested, and RT-qPCR was performed. The heat map shows the relative expression of genes involved in metabolism, normalized to the untransduced Treg expression level. The depicted results were generated from a combination of four independent experiments. G6PD: glucose-6-phosphate dehydrogenase; HGPRT: hypoxanthine-guanine phosphoribosyltransferase; SLC1A5: solute carrier family 1, member 5; SLC38A1: solute carrier family 38, member 1; TPI: triosephosphate isomerase. B. ASCT2 expression measured by flow cytometry on day 16 of culture. The ratio of fluorescence intensity (RFI) was calculated as follows: the geometric mean fluorescence (MFI) of either CD28 or 4-1BB CAR-Tregs divided by the MFI of untransduced Tregs. Each replicate is represented. N=7 donors from 7 independent experiments. Wilcoxon matched-pairs signed rank test, *p<0.05. C-D. The frequency of phosphoS6-positive cells among living cells determined by flow cytometry on day 16 of culture. C. Representative histograms of phosphoS6 staining. D. The mean +/- SEM of n=5 donors from five independent experiments is shown. E. qPCR analysis of cells lysed on day 16 showing the percent demethylation of CNS2 within the *FOXP3* locus, known as the Treg-specific demethylated region (TSDR). n=5 donors from five independent experiments. F-I. Flow cytometry analysis of HELIOS and FOXP3 expression in live cells during culture. F. Representative contour plots for day 16. G. Frequency of FOXP3-positive cells on day 16 from n=12 donors (ten independent experiments). H. Frequency of double-positive FOXP3+ HELIOS+ cells on day 16 from n=12 donors (ten independent experiments). H. Frequency of double-negative (HELIOS and FOXP3) live cells on days 10, 16 and 21 of culture, as assessed by flow cytometry. The results for n=4-6 donors from at least four independent experiments are represented. The mean +/- SEM is represented. Mann-Whitney tests. *p<0.05, **p<0.01.

mTORC1 is crucial in supporting the proliferative capacities of Tregs (Gerriets et al., 2016; Procaccini et al., 2016). Hence, we studied mTORC1 activity, as assessed by the phosphorylation of S6 (pS6) in 4-1BB CAR-Tregs and CD28 CAR-Tregs on day 16 of culture. In contrast to CD28 CAR-Tregs and untransduced Tregs, 4-1BB CAR-Tregs displayed high expression of pS6 five days after TCR stimulation.

Although mTOR plays roles in Tregs (Gerriets et al., 2016; Procaccini et al., 2016) and functional differentiation (Zeng et al., 2013), its overactivation can reprogram Tregs toward Tconvs. We thus analyzed the stability of the Treg lineage according to the CAR CSD. To this end, we studied the demethylation status of the Treg-specific demethylated region (TSDR) and found that CAR-Tregs retained a high degree of TSDR demethylation on day 16, irrespective of the CSD (Figure 4E). Moreover, the expression of FOXP3 and HELIOS was assessed throughout the culture period. On day 16 (Figure 4F-H), the proportions of FOXP3+ (Figure 4G) and double-positive FOXP3+ HELIOS+ (Figure 4H) Tregs were similar between 4-1BB CAR-Tregs and CD28 CAR-Tregs and were much greater than those of Tconvs. We concluded that CAR-Treg manufacturing, including lentiviral transduction, CAR tonic signaling and TCR-driven two-week culture, did not unduly disrupt Treg stability, regardless of the CSD, at early time points. However, a double-negative FOXP3-HELIOS-population was readily detected as early as day 16 in 4-1BB CAR-Tregs and subsequently expanded in all conditions (Figure 4I). This finding raised the concern that prolonged culture and extensive proliferation, especially that driven by 4-1BB tonic signaling, could promote Treg instability.

### Combination of rapamycin and vitamin C abates overactivation due to 4-1BB tonic signaling and improves CAR-Treg stability

Uncontrolled mTOR pathway activation and related hyperglycolytic and hyperglutaminolytic states promote Treg destabilization (Gerriets et al., 2016; Huynh et al., 2015; Xu et al., 2017). We thus hypothesized that mTOR inhibition could mitigate metabolic reprogramming and prevent 4-1BB CAR-Treg instability (Battaglia et al., 2006). Moreover, glutamine metabolites inhibit Ten Eleven Translocation (TET) enzymes that demethylate the *FOXP3* gene (Xu et al., 2017), whereas vitamin C, a potent activator of TET enzymes, promotes Treg stability (Lu et al., 2017; Sasidharan Nair et al., 2016). We next wondered whether culturing 4-1BB CAR-Tregs with a Treg-supporting cocktail, including the mTOR inhibitor rapamycin and the epigenetic modifier vitamin C, would improve the maintenance of the 4-1BB CAR-Treg lineage. Rapamycin and vitamin C were added to the culture medium at the time of restimulation (day 11) and compared with complete medium alone. As expected, the addition of rapamycin and vitamin C abated the activation of the glycolytic and glutaminolytic pathways (Figure 5A) and strongly inhibited mTORC1 activity (Figure 5B-C). In line with these findings, the expansion rate of 4-1BB CAR-Tregs decreased to a level comparable to those of CD28 CAR-Tregs and untransduced Tregs (Figure 5D). As a positive effect, reduced proliferation correlated with greater 4-1BB CAR-Treg stability, according to phenotypic (Figure 5E) and epigenetic assessments (Figure 5F). For subsequent *in vivo* experiments, the Treg-supporting cocktail was added to the culture medium from days 10 to 18 for 4- 1BB CAR-Tregs, whereas unmodified medium was used for other Treg populations.

**Figure 5:**
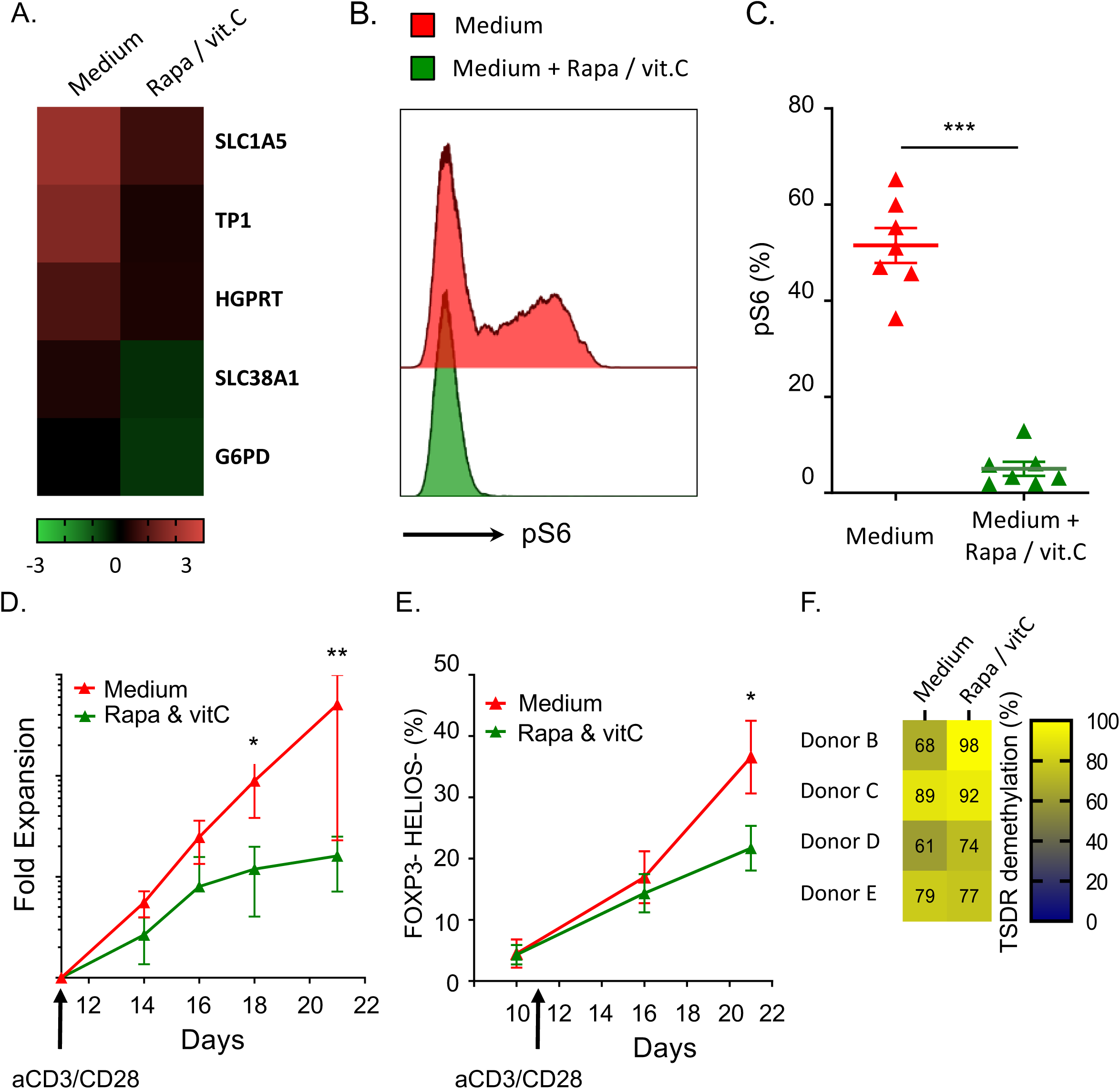
A rapamycin/vitamin C cocktail curbs 4-1BB-induced CAR-Treg overactivation. A. Heat map representation of the relative expression of genes involved in metabolism analyzed on day 16 by qRT-PCR for 4-1BB CAR-Tregs with or without rapamycin and vitamin C treatment, normalized to the untransduced Treg expression level. The depicted results were generated from a combination of four independent experiments. G6PD: glucose-6-phosphate dehydrogenase; HGPRT: hypoxanthine-guanine phosphoribosyltransferase; SLC1A5: solute carrier family 1, member 5; SLC38A1: solute carrier family 38, member 1; TPI: triosephosphate isomerase. B-C. From day 9 to d 16 of culture, Tregs were harvested, and the percentage of ps6-positive single cells was determined by flow cytometry at the indicated time points. B. Representative histogram C. The results of n=3 donors from at least four independent experiments are represented. The median is represented. D. Fold expansion of CD28 or 4-1BB CAR-Tregs from restimulation on day 11 to day 21. The mean+/- SEM is represented. n=4 donors, pooled from at least 4 independent experiments. E. The percentage of double-negative (HELIOS and FOXP3) live cells was determined on days 10, 16 and 21 of culture by flow cytometry. The results of n=4-6 donors from at least four independent experiments are represented. F. Percentage of 4-1BB CAR-Tregs with Treg-specific demethylated region (TSDR) demethylation on day 16 of culture with or without rapamycin and vitamin C beginning on day 11. The mean +/- SEM is represented. Mann-Whitney tests. *p<0.05, **p<0.01, ***p<0.001.

### Both 4-1BB CAR-Tregs and CD28 HLA-A2 CAR-Tregs are efficient in suppressing xenogeneic graft-versus-host disease (xenoGVHD)

To assess the suppressive capacities of CAR-Tregs *in vivo* in an antigen-specific manner, we used a xenogeneic GVHD model based on the transfer of thawed HLA-A2-negative peripheral blood mononuclear cells (PBMCs) into busulfan-conditioned NSG mice. Different doses of hPBMCs were tested (5, 10 or 20 x 10^6^ hPBMCs). The onset of GVHD symptoms occurred after 10 days, while the GVHD clinical score peaked at 40 days (Figure Supp 2A). The median survival times were 24, 28 and 38 days for 20, 10 and 5 x 10^6^ hPBMCs, respectively (Figure Supp 2B). The mean percentage (±SD) of human chimerism in the blood on day 10 was measured to be 2.4±2.1, 21.6±17.5, and 41.6±21.5% in the animals transferred with 5, 10 or 20 x 10^6^ hPBMCs, respectively, demonstrating a dose-dependent effect (Figure supp 2C). Of note, the levels of human IFN-γ in the plasma were significantly correlated with human cell engraftment (Spearman r= 0.74; Figure Supp 2D). The main target organs included the lungs and liver, which were massively infiltrated by leukocytes (Figure Supp 2E).

Next, we wanted to compare HLA-A2 CAR-Tregs with control polyclonal Tregs (transduced with an EGFRt-expressing vector, referred to as MOCK-Tregs) in regard to xenoGVHD prevention efficacy. Notably, 5×10^6^ hPBMCs and 5×10^6^ Tregs were injected at the same time through different intravenous routes (retroorbital sinus and tail vein). This precaution stemmed from a previous finding showing that mixing HLA-A2+ PBMCs and HLA-A2-targeted CAR-Tregs before infusion impedes CAR-Treg homing (not shown). CAR-Tregs and MOCK-Tregs were cotransduced with a lentivirus encoding luciferase and the mCherry reporter gene to allow *in vivo* tracking (Figure 6A). Regarding survival, all the mice transferred with PBMCs alone died within 60 days, as expected, whereas polyclonal MOCK-Tregs slightly delayed the disease course (median survival: 32 days for PBMCs vs 43 days for MOCK-Tregs, p=0.091). In contrast, both HLA-A2-specific 4-1BB CAR-Tregs and HLA-A2-specific CD28 CAR-Tregs produced significant protection against xenoGVHD and related death (Figure 6B). Moreover, circulating hIFN-γ levels were significantly lower in mice that received either CD28 CAR-Tregs or 4-1BB CAR-Tregs than in animals transferred with PBMCs alone (Figure 6C). These results agreed with those for human chimerism in the blood, which was strongly inhibited by cotransfer of CAR-Tregs (Figure 6D). Strikingly, circulating CAR-Tregs were readily detected on day 24 (Figure 6D), far later than in previous reports (Dawson et al., 2020; Nowak et al., 2018). In our hands, a sizeable population of CAR-Tregs could be detected in the blood (median: 4.4 [0.2-9.1]% of circulating cells) as late as day 60-62 for CD28 CAR-Tregs, which was when mice were sacrificed (Figure Supp 3A). In contrast, polyclonal MOCK-Tregs vanished very rapidly from the peripheral blood (Figure 6D) and were barely detectable after day 10. In line with these results, we performed histologic analysis of both the liver and lungs at the time of sacrifice, and we observed a significant decrease in the histologic score of xenoGVHD in mice that received CD28 CAR-Tregs or 4-1BB CAR-Tregs compared to those treated with MOCK-Tregs (Figure Supp 3B-C). We concluded that both 4-1BB CAR-Tregs and CD28 CAR-Tregs were efficient at preventing xenogeneic acute GVHD through early inhibition of hPBMC expansion.

**Figure 6:**
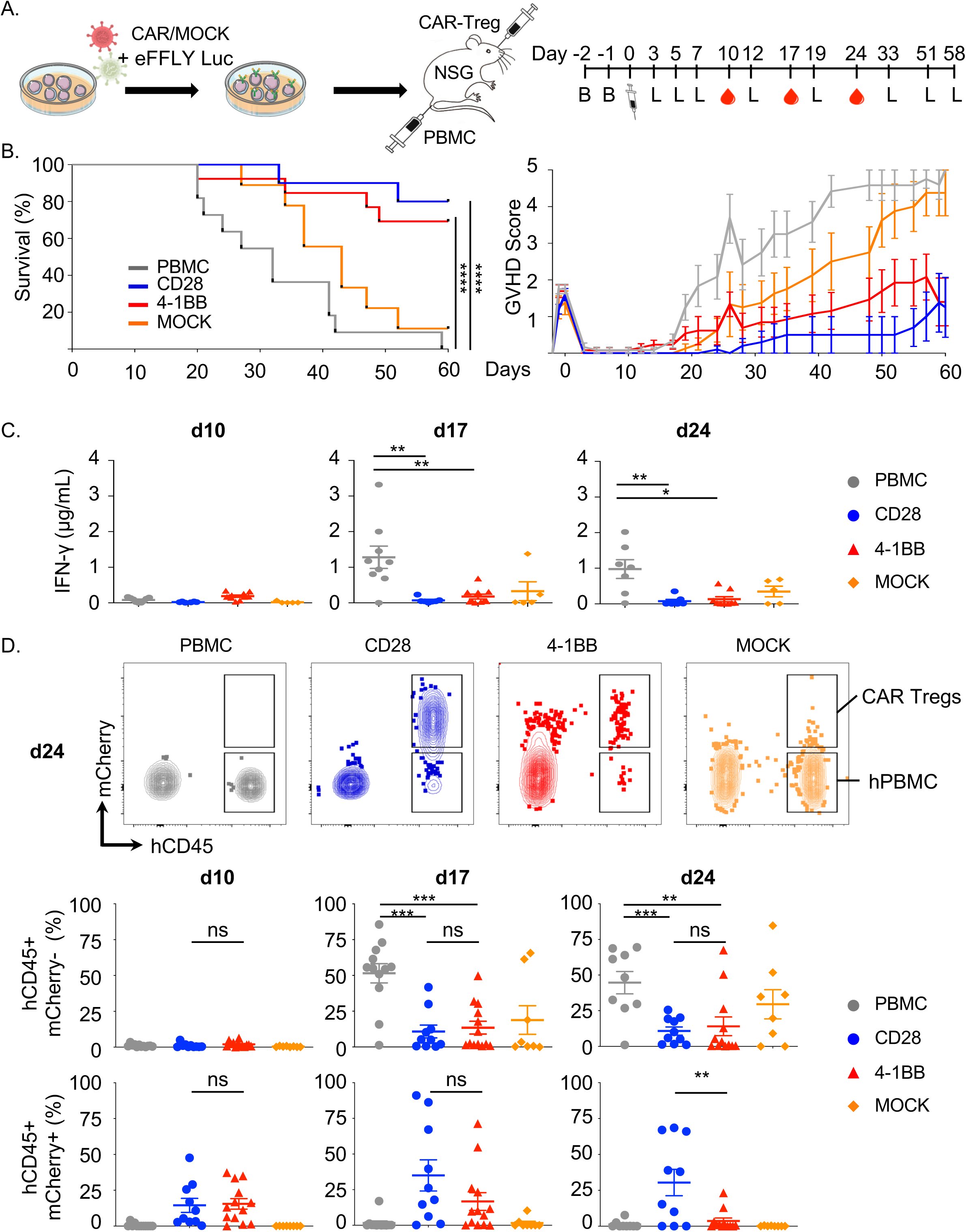
Both CD28 CAR-Tregs and 4-1BB CAR-Tregs efficiently suppress acute xenoGVHD. Eight- to 12-week-old male NSG mice were conditioned with busulfan on day -2 and day -1 before IV injection of 5.10^6^ HLA-A2+ PBMCs followed by retro-orbital injection of 5×10^6^ of the indicated type of CAR-Tregs (n=11, 13, 13 and 10 for PBMCs, CD28 CAR-Tregs, 4-1BB CAR-Tregs and MOCK-Tregs, respectively). Mice were weighed and scored for GVHD three times a week and bled weekly for flow cytometry analysis. A. Schematic design of the experiment. B. Survival curves (left panel) and GVHD scores (right panel). Log-rank Mantel-Cox tests comparing the indicated groups. *p<0.05. C. Ten, 17 and 24 days post-cell injection, the concentration of human IFN gamma in the plasma was measured by a cytometry bead array. D. Representative contour plots of circulating cells, including murine cells (hCD45), CAR/MOCK-Tregs (mCherry+, hCD45+), and xenoreactive PBMCs (mCherry-, hCD45+), across the different groups (upper panels). Frequencies of hCD45+ mCherry-cells (middle panels) and CAR-expressing (mCherry+) cells (lower panels) among the total circulating cells. Abbreviations: B, busulfan; L, bioluminescence

### 4-1BB limits the *in vivo* persistence of CAR-Tregs

Luciferase-expressing CAR-Tregs were found to rapidly traffic through the lymph nodes and subsequently accumulate in the spleen (Figure 7A). Using an *In Vivo* Imaging System (IVIS) Spectrum optical/CT coregistration system, we were able to not only map CAR-Tregs reliably in organs but also provide accurate quantification of cells in a 3D-specific region of interest (Figure 7B). In line with the blood data, we found that MOCK-Tregs rapidly disappeared, whereas HLA-A2-specific CAR-Tregs exhibited long-term persistence *in vivo*. Despite peaks of comparable magnitude, CD28 CAR-Tregs were detected in the spleen longer than 4-1BB CAR-Tregs throughout the follow-up period after day 12. Indeed, the number of splenic CD28 CAR-Tregs plateaued and remained constant from day 12 to day 58, whereas that of splenic 4-1BB CAR-Tregs slowly decreased over time until reaching a significant decrease on day 58 (median absolute cell count: 2.0×10^4^ vs 4.4×10^6^ for 4- 1BB and CD28 CAR-Tregs, respectively, p=0.0127; Figure 7B, right panel). Hence, the *in vivo* persistence of 4-1BB CAR-Tregs was significantly decreased in the long term compared to that of CD28 CAR-Tregs.

**Figure 7:**
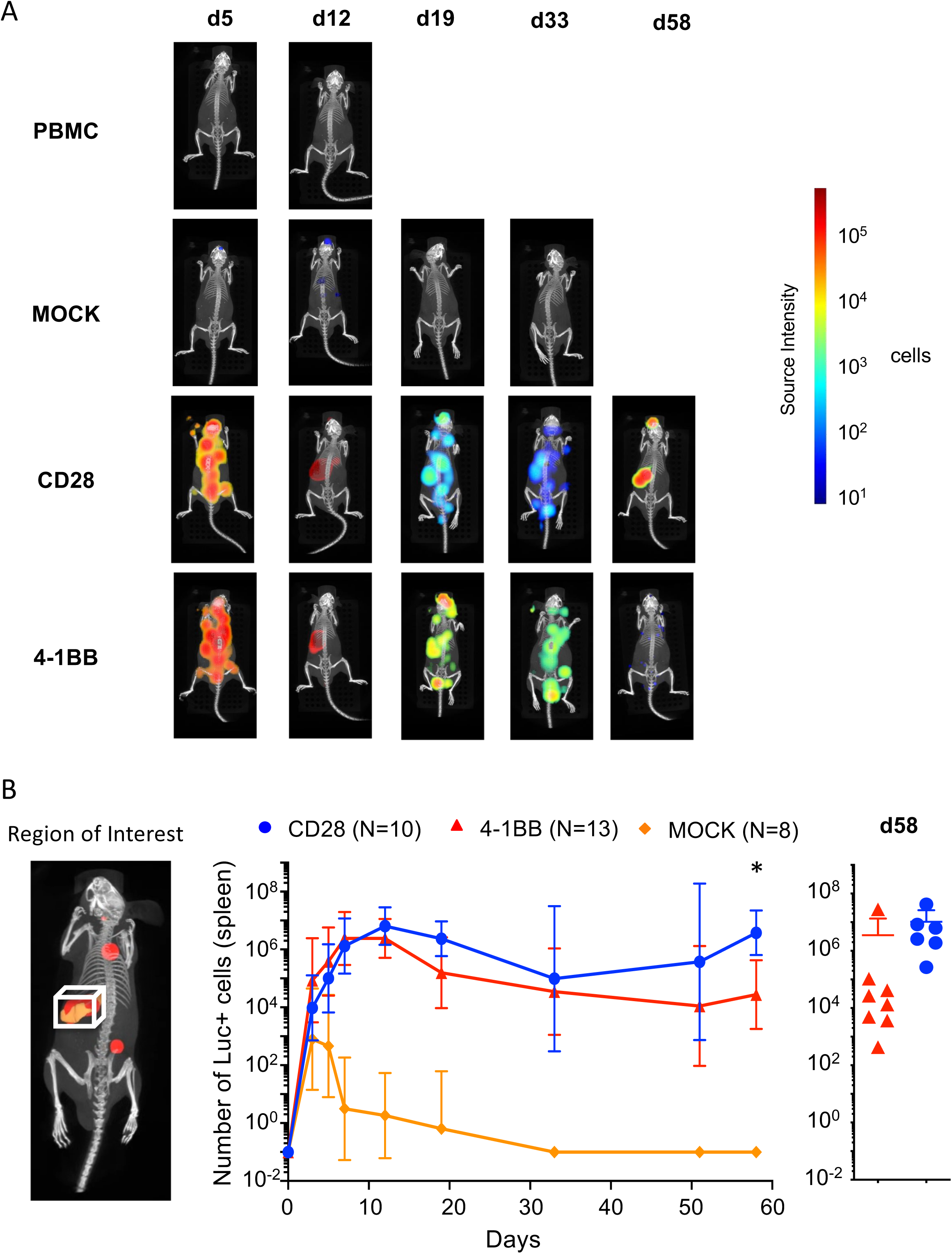
CD28-CAR-Tregs persisted for longer than 4-1BB-CAR-Tregs. A. 3D bioluminescence imaging tomography (DLIT) of recipient mice was performed as described in the Methods. B. A region of interest (ROI) corresponding to the spleen was determined by coregistration with the Automatic Mouse Atlas, and the absolute cell number of luciferase-positive (Luc+) cells was calculated with *in vitro* calibration with corresponding cells. Kinetics of the absolute numbers of polyclonal MOCK-Tregs and CD28 or 4-1BB CAR-Tregs in the spleen. At least three independent experiments and three different donors are represented. The number of CAR-Tregs in the spleen of surviving animals on day 58 was calculated according to the CAR CSD (right panel). The mean +/- SEM is represented. Mann-Whitney tests. *p<0.05, **p<0.01, ***p<0.001.

### 4-1BB induces global hyporesponsiveness over long-term culture *in vitro*

We wondered whether the shorter *in vivo* persistence of CAR-Tregs could be associated with tonic signaling-related functional impairment, similar to that seen in effector T cells (Ajina and Maher, 2018). Although Treg exhaustion is still ill-defined, we hypothesized that strong tonic signaling could produce hyporesponsiveness in 4-1BB CAR-Tregs following restimulation. To address this issue, cultured CAR-Tregs and untransduced Tregs, which were collected after 16 days of culture, were stimulated overnight with either CD3/CD28 beads or plate-bound HLA-A2 pentamers. Strikingly, 4-1BB CAR-Tregs failed to properly upregulate the expression of activation markers, such as CD69 (Figure 8A) and 4-1BB (Figure 8B), especially after CAR stimulation. The hyporesponsiveness of the 4-1BB CAR-Tregs was in sharp contrast to the full ability of CD28 CAR-Tregs and polyclonal Tregs to respond to antigen receptor stimulation. Of note, pretreatment with an mTOR inhibitor failed to restore sensitivity to CAR stimulation. Furthermore, both CD25 and HLA-DR seemed to be highly and continuously expressed by 4-1BB CAR-Tregs, regardless of the culture conditions (with or without the mTOR inhibitor) (Figure 8C and D). Taken together, these findings suggest that 4-1BB CAR tonic signaling promotes accelerated dysfunction in Tregs, consistent with the lack of ability to be restimulated through antigen receptors.

**Figure 8:**
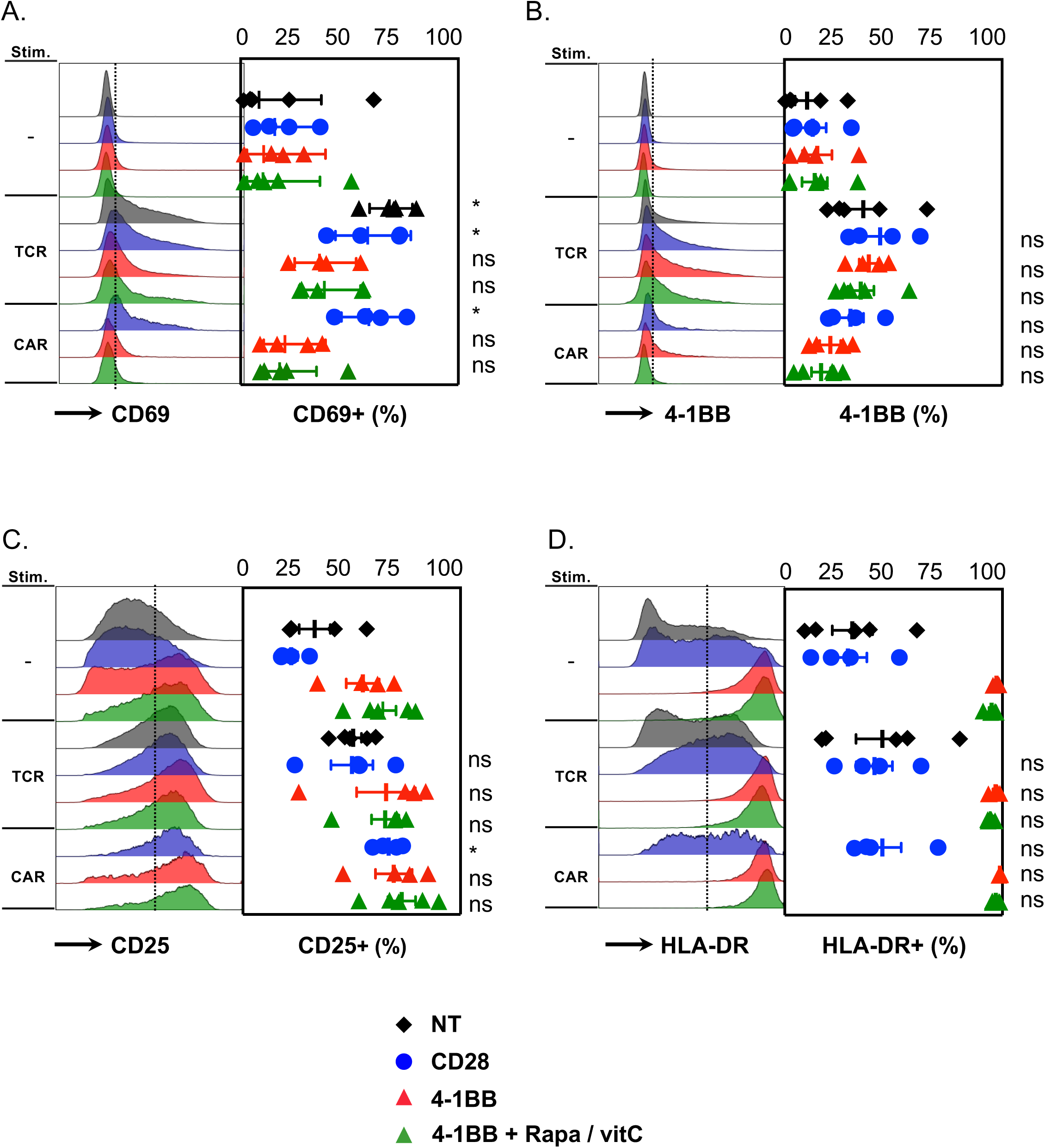
4-1BB CSD drives CAR-Treg hyporesponsiveness. A-D. Human Tregs were transduced and expanded as shown in Figure 1C. On day 16, cells were stimulated overnight with either plate-bound HLA-A2 pentamers or CD3/CD28 beads before FACS analysis. Representative histograms of each activation marker (left panels) and the frequencies of positive cells (right panels) before and after stimulation are shown. n=5 donors, pooled from at least 4 independent experiments. The mean +/- SEM is represented. Mann-Whitney tests were performed to compare unstimulated cells and stimulated (either via the TCR or CAR) cells transduced with the same CAR construct. *p<0.05.

## DISCUSSION

Adoptive regulatory cell therapy represents a promising strategy to promote operational tolerance in solid organ transplantation. Although substantial efforts have been made to understand the effects of the CSD in CAR-T cells, this important issue has only recently started to be investigated in Tregs (Dawson et al., 2020; Nowak et al., 2018). In this study, we used HLA-A2-targeted CARs in a transplant model to demonstrate that the CD28 CSD more strongly abates the negative impact of CAR tonic signaling on Treg lineage stability and function than does the 4-1BB CSD.

In conventional T cells, CAR molecules frequently produce antigen-independent tonic signaling, commonly promoted by the high cell-surface density and intrinsic self-aggregating properties of CARs (Ajina and Maher, 2018). However, in our settings, the extracellular scFv, lentiviral VCN and CAR cell-surface density were comparable between the CD28 and 4-1BB CARs. We thus inferred that the more deleterious tonic signaling associated with 4-1BB, compared to CD28, could neither be ascribed to a specific characteristic of the scFv nor to greater transgene expression. In contrast, our data supported an intrinsic role for the 4-1BB CSD in enhancing the negative impact of tonic signaling on Treg biology. Previous studies showed that 4-1BB CAR-Treg proliferation was greater than that of CD28 CAR-Tregs following CAR stimulation (Dawson et al., 2020; Nowak et al., 2018). We extended this finding by showing the interference between 4-1BB tonic signaling and pathways downstream of TCR activation, resulting in heightened proliferation and increased expansion of 4-1BB CAR-Tregs upon CD3/CD28 stimulation.

We observed that both CAR-Treg subsets consistently exhibited unique phosphorylation patterns for downstream signaling proteins, which were present shortly after TCR stimulation, that markedly differed from that of untransduced Tregs. These findings underscore the presence of tonic signaling triggered by both CAR constructs but the involvement of different pathways. Notably, the 4-1BB CSD induced much stronger activation of the ERK1/2 MAPK pathway than did the CD28 CSD. The p38-MAPK signaling pathway was found to be instrumental in TNFR2-driven Treg proliferation (He et al., 2018) and in the transduction of 4-1BB-induced activation in CD8+ T cells (Menk et al., 2018). However, our study is the first to show a link between 4-1BB-induced Treg proliferation and MAPK activation. On the other hand, epigenetic marks along with constitutive expression of dual-specificity phosphatase 4 (DUSP4), a potent regulator of the MAPK superfamily, are highly conserved features of Tregs across species (Ferraro et al., 2014; Arvey et al., 2015). Furthermore, DUSP4-dependent control of the MAPK pathway was found to be critical for the maintenance of Treg function (Sugimoto and Liu, 2013). Together, these results suggest that unduly activating MAPK through 4-1BB tonic signaling in Tregs could impede Treg function. Whether 4-1BB-induced MAPK overactivation can lead to increased tyrosine hydroxylase expression in Tregs (Cosentino et al., 2007), resulting in enhanced proliferation and decreased function, has yet to be investigated.

Surprisingly, the impact of CARs containing the CD28 CSD or the 4-1BB CSD on T cell fate greatly varies between Tregs and Tconvs. In Tconvs, the 4-1BB CSD promotes mitochondrial biogenesis and an oxidative metabolic program that supports long-lived central memory T cells (Kawalekar et al., 2016; Long et al., 2015; Milone et al., 2009; Zhao et al., 2015), whereas the CD28 CSD enhances an aerobic glycolytic burst, which accompanies differentiation into effector memory T cells (Kawalekar et al., 2016). In contrast, our results showed that 4-1BB CAR-Tregs promptly progressed to a highly differentiated CD45RA-CCR7- effector memory phenotype, whereas CD28 CAR-Tregs better maintained CCR7 expression. Furthermore, 4-1BB CAR-Tregs exhibited higher levels of function-associated receptors and activation molecules, such as CD39, HLA-DR and 4-1BB, consistent with the phenotype of effector Tregs. This finding is reminiscent of data in mice showing the key role of the TNFR superfamily in the generation of effector Tregs (Vasanthakumar et al., 2017).

There is growing evidence establishing the key role of metabolism in regulating immune cell fate decisions (Chi, 2012; Galgani et al., 2016). In Tregs, although mTORC1 activity and the ensuing anabolic state are required for Treg proliferation and acquisition of full suppressive function (Zeng et al., 2013; Procaccini et al., 2016), it is also well established that the PI3K/Akt/mTOR pathway must be tightly regulated through a feedback loop involving phosphatase and tensin homolog (PTEN) to enforce Treg stability (Huynh et al., 2015; Newton et al., 2016). In fact, unregulated activity of mTOR (Huynh et al., 2015) and related metabolic programs, including glycolysis (Gerriets et al., 2016) and glutaminolysis (Xu et al., 2017), can interfere with *FOXP3* epigenetics and promote Treg lineage instability. Hence, mTORC1 is increasingly recognized as the fulcrum of Treg regulation, balancing proliferative and suppressive capacities (Gerriets et al., 2016; Newton et al., 2016). In this respect, we found that 4-1BB tonic signaling enhanced mTOR activity (both mTORC1 and mTORC2) and induced glutamine transporter expression. The metabolic changes observed in 4-1BB CAR-Tregs were accompanied by a higher rate of HELIOS and FOXP3 loss than those in CD28 CAR-Tregs. HELIOS plays a key role in IL2-dependent Treg stability (Kim et al., 2015), and its expression is associated with the most stable Treg subset in humans (Bin Dhuban et al., 2015). The addition of a Treg-supporting cocktail (including vitamin C, a Treg epigenetic stabilizer (Yue et al., 2016), and rapamycin, an mTOR inhibitor) to cultured 4-1BB CAR-Tregs reduced FOXP3 loss and better maintained TSDR demethylation, but these effects came at the expense of the fold expansion in culture.

Tonic CAR signaling can induce early exhaustion in CAR-T cells, producing a shortened lifespan after transfer and reduced antitumor efficacy (Long et al., 2015). Although the term “exhaustion” is broadly accepted to refer to effector T cells with progressive functional impairment and reduced proliferative potential along with an accumulation of inhibitory receptors (Blank et al., 2019), there is no equivalent state for Tregs. Given the constitutive expression of a number of inhibitory receptors by Tregs, including CTL-4, LAG3, TIM-3, and TIGIT, the study of these receptors is of little value. However, emerging data suggest that chronically stimulated human Tregs may also progress toward dysfunction associated with TSDR remethylation and increased IFN-γ secretion (Lowther et al., 2016). Similarly, in our settings, we found that enhanced tonic signaling in CAR-Tregs could lead to reduced Treg survival *in vivo* and an altered ability to be restimulated through the CAR. Strikingly, although CD28 was found to worsen the process of tonic signaling-induced exhaustion in Tconvs (Long et al., 2015), our data showed the opposite in Tregs. The respective effects of the 4-1BB and CD28 CSDs on Tregs are a mirror image of those on Tconvs.

Despite the abovementioned differences between the two CSDs, both CD28 CAR-Tregs and 4-1BB CAR-Tregs maintained the overall features of Tregs, including limited inflammatory cytokine production, reduced cytotoxicity, TSDR demethylation and *in vivo* suppressive activity. These findings conflict with two recent studies reporting the lack of *in vivo* suppressive function of 4-1BB CAR-Tregs (Boroughs et al., 2019; Dawson et al., 2020). The reason for these discrepancies is not clear. Remarkably, 4-1BB CAR-Tregs potently suppressed xenoGVHD despite a reduced ability to be stimulated through the CAR. This finding is reminiscent of a previous report showing *in vivo* antigen-specific suppression mediated by HLA-A2-targeted CAR-Tregs, even in the absence of signaling domains in the CAR structure (Boardman et al., 2017). Together, these data suggest that the localization of preactivated CAR-Tregs through antigen binding could be critical for their protective effect, even if the CAR fails to be stimulated.

Finally, we were struck by the much longer persistence of circulating and lymphoid organ-resident CAR-Tregs in our xenoGVHD model than that reported in previous studies. It was previously reported that circulating CAR-Tregs were hardly detected after 7 days (Dawson et al., 2019, 2020). In contrast, in our hands, CAR-Tregs were readily detected in the peripheral blood on day 10 regardless of the CSD and could be easily found as late as day 60 in most animals transferred with CD28 CAR-Tregs. A simple explanation for these differences could be that we transferred CAR-Tregs and HLA-A2+ PBMCs through different routes (retroorbital sinus and tail vein) to avoid cell clustering and early trapping in the lungs following *in vivo* transfer. In fact, immediate encounter of CD19+ target B cells in the blood by CD19-specific CAR-T cells was found to trap the CAR-T cells in the lungs and reduce their access to the lymphoid organs (Cazaux et al., 2019; Iwano et al., 2018).

In conclusion, we found that constitutive CAR signaling might induce an imbalance between proliferation and suppressive function, whose equilibrium in Tregs is normally controlled by finely tuned TCR signal transduction pathways and metabolic checkpoints. Hence, CAR tonic signaling could interfere with the fate of CAR-engineered Tregs, similar to the effects on Tconvs. Notably, the transduction pathways involved in downstream CAR activation varied according to the CSD and resulted in different metabolic and activation/differentiation patterns. More importantly, unlike in Tconvs, we found that the CD28 CSD but not the 4-1BB CSD was protective against tonic signaling-induced dysfunction and destabilization.

## MATERIALS & METHODS

### scFv affinity and specificity

The affinity and specificity of different scFvs for HLA-A2-bound beads were assessed by a collaborating team using a Luminex multiplex single antigen bead assay (LABScreen® Single Antigen). A cell-based assay was used to confirm HLA allele specificity. Tagged low- and high-affinity scFvs at three different concentrations, 0.05 µg/mL, 0.5 µg/mL, and 5 µg/mL, were incubated with PBMCs from HLA-A2+ and HLA-A2-/A28- donors. The MFIs of the tagged scFvs were measured by FACS analysis.

### CAR construct design

The selected scFv was fused to a stalk region from the human CD8α hinge, the transmembrane domain from human CD8, the CD28 or 4-1BB CSD, and the human CD3ζ signaling domain. EGFRt was cloned downstream of T2A at the second gene position to serve as a reporter (Figure Supp 1D).

### Cell source and purification of human Tregs

Healthy donor PBMCs were obtained from the Etablissement Français du Sang. PBMCs were typed based on the expression or absence of HLA-A2/A28 molecules, as assessed by anti-HLA-A2/A28 antibody (OneLambda) staining evaluated by FACS analysis. CD4+ T cells were enriched from HLA-A2- donor PBMCs using an EasySep CD4+ Enrichment kit (Stem Cell®). Naïve regulatory T cells (nTregs), which were defined as CD4+ CD25++ CD45RA+ CD127low, were sorted using a FACSAria II (BD Biosciences). In parallel, we sorted CD4+ CD45RA+ CD25-Tconv cells from the same donor as controls. Sorted T cells were stimulated with Dynabeads® Human T-Activator CD3/CD28 (Thermo Fisher Scientific) (ratio 1:1) in X-VIVO® 20 medium containing 10% human serum AB (Biowest) and 1000 UI/mL human IL-2 (Proleukin, Novartis).

HLA-typed CD3-depleted splenocytes were isolated from spleens collected from deceased organ donors through a collaboration with the Regional Histocompatibility Laboratory and National Biomedicine Agency after local ethics committee approval (Project n°: 2016-12-01; approved on December 5, 2016).

### T cell transduction and expansion

Two days after activation, Tregs were transduced with the different CAR constructs at a multiplicity of infection (MOI) of 30 viruses per cell. Prostaglandin E2 (PGE2) (Cayman Chemical, 10 µM) and a transduction adjuvant (Lentiboost, Sirion Biotech, 0.25 mg/mL) were added for a 6 h of transduction time. On day 7 posttransduction, CD4+ EGFRt+ cells were sorted using a FACSAria II and then restimulated with CD3/CD28 Dynabeads®. T cells were expanded and cultured in complete medium supplemented with IL-2 (1000 UI/mL) for 7 to 10 days for *in vitro* experiments.

### Flow cytometry

Surface staining was performed with an HLA-A2 pentamer (Proimmune) and the monoclonal antibodies listed in Supplemental Table 2. For CAR surface expression assessment, biotinylated protein L (Thermo Fisher Scientific) was used and bound to fluorochrome-conjugated streptavidin (BD Biosciences). For ASCT2 expression evaluation, a receptor-binding domain (Metafora) was preincubated for 20 minutes at 37°C with cells, and an anti-mouse secondary antibody was used. pS6 staining was performed using a PerFix EXPOSE kit (Beckman Coulter). For evaluation of intracellular targets, cells were stained with a fixable viability dye (FVD, Thermo Fisher Scientific) before fixation, permeabilization and staining using a FOXP3/transcription factor staining buffer set (Thermo Fisher Scientific) according to the manufacturer’s instructions. Intracellular staining was performed for FOXP3, HELIOS (Thermo Fisher Scientific), Granzyme B and Ki67 (Thermo Fisher Scientific). Cells were analyzed using a Fortessa X-20 cytometer (BD Biosciences). Data were analyzed using Kaluza 2.1 (Beckman Coulter) or FlowJo 10.6.1 (BD Biosciences). For t-SNE mapping, the R Console plugin (https://www.beckman.fr/flow-cytometry/software/kaluza/r-console) was used to analyze flow cytometry data in R within the Kaluza Analysis framework.

### Activation assay

Ninety-six-well flat-bottom plates were coated overnight at 4°C or for 3 h at 37°C with an HLA-A2 pentamer at 5 µg/mL in PBS. The plates were washed with cold PBS the next day or 3 hours later. Cells were separated from CD3/CD28 Dynabeads® 24 h prior to seeding at 100,000 cells/well.

### Immunoblotting

T cells (3-5 x 10^6^) were lysed in 1% NP-40, 50 mM Tris pH 8, 150 mM NaCl, 20 mM EDTA, 1 mM Na3VO4, 1 mM NaF, a complete protease inhibitor cocktail (Roche), and anti-phosphatase cocktails 2 and 3 (Sigma-Aldrich). Protein concentrations were quantitated with a BCA assay (Bio-Rad). Eighty micrograms of protein was separated by SDS-PAGE and transferred to PVDF membranes (Millipore). The membranes were blocked with milk or BSA for 1 h before incubation with primary antibodies for 1.5 h. The following mAbs and rabbit polyclonal antibodies were used for immunoblotting: anti-PLC-γ1 (clone D9H10), anti-phosphorylated PLC-γ1 (clone D6M9S), anti-phosphorylated ERK1/2 (clone D13.14.4E), anti-ERK1/2 (clone 137F5), anti-phosphorylated Akt (clone D9E), anti-Akt (clone C67E7), anti-Ku80 (clone C48E7), and anti-phosphorylated tyrosine (clone Tyr-100) purchased from Cell Signaling Technology. The membranes were then washed and incubated with anti-mouse or anti-rabbit HRP-conjugated secondary antibodies from Cell Signaling Technology and GE Healthcare, respectively. Pierce ECL western blotting substrate was used for detection as previously described (Hauck et al., 2012; Latour et al., 2001). Quantification was performed using ImageJ software according to the standard gel signal measurement method.

### Cytokine analysis

Plasma or culture supernatants were collected at the indicated times, and cytokine production (TNF-α, IFN-γ, IL-2, IL-4, IL-6, IL-10, and IL-17A) was determined using the Cytometric Bead Array Th1/Th2/Th17 Kit (BD Biosciences) according to the manufacturer’s instructions.

### RNA extraction and quantitative real-time polymerase chain reaction (qRT-PCR)

mRNA was extracted using the RNeasy Mini Kit® (Qiagen) according to the manufacturer’s protocol and subjected to reverse transcription (High Capacity RNA-to-cDNA Master Mix (Applied Biosystems™)). mRNA expression levels were assessed using Fast SYBR™ Green Master Mix (Applied Biosystems™) with a Viia7 thermocycler (Thermo Fisher Scientific). The fold-change for each tested gene was normalized to that of the housekeeping gene ribosomal protein L13A (RPL13A). The relative expression of each gene was calculated using the 2−ΔΔCT method (Livak and Schmittgen, 2001). Consequently, the expression level of a given gene in control samples (untransduced Tregs) determined using the 2−ΔΔCT method was set to 1. The results were visualized as log2(relative expression) in heat maps. The primer sequences are listed in Supplementary Table 1.

### VCN assessment

Genomic DNA was isolated from transduced cells using a Genomic DNA Purification kit (Qiagen). gDNA was digested with HindIII HF (New England Biolabs, Evry, France) in a total reaction mixture of 6 µl at 37°C for 30 minutes. Droplet Digital™ PCR (ddPCR) was performed with TaqMan probes designed to detect a lentiviral sequence (Psi) (Bio-Rad) and a sequence in the human genome (Albumin) (Thermo Fisher Scientific). The final reaction mixture contained ddPCR Mastermix (Bio-Rad), forward and reverse primers, probe solutions, and digested gDNA. The sample mixture was transferred to a DG8 cartridge and placed into the QX100 droplet generator (Bio-Rad). Sample droplets were transferred to a 96-well PCR plate, and ddPCR was performed using a SimpliAmp thermal cycler (Thermo Fisher Scientific). The plate was analyzed using a QX200 droplet reader (Bio-Rad). Using the manufacturer’s QuantaSoft software (Bio-Rad), the concentration of the target amplicon per unit volume of input for each sample was estimated for both Psi and the Albumin reference gene. The VCN was determined for each sample as follows: VCN = 2 x Psi signal/Albumin signal.

### TSDR DNA Methylation Analysis

Regulatory T cell-specific demethylation region (TSDR) DNA methylation analysis was performed as previously described (Wieczorek et al., 2009) using genomic DNA isolated from freshly sorted or expanded Treg cells using the QIAamp DNA Mini Kit (Qiagen). A minimum of 60 ng bisulfite-treated (EpiTect; Qiagen) genomic DNA was used for qRT-PCR to quantify *∑* TSDR demethylation. Real-time PCR was performed in a final reaction volume of 20 µL containing 10 µL FastStart universal probe master (Roche Diagnostics), 50 ng/µL lambda DNA (New England Biolabs), 5 pmol/µL methylation or nonmethylation-specific probe, 30 pmol/µL methylation or nonmethylation-specific primers, and 60 ng bisulfite-treated DNA or a corresponding amount of plasmid standard. Samples were analyzed in triplicate on an ABI 7500 cycler (Thermo Fisher Scientific), and results are reported as % T cells with a demethylated TSDR region.

### XenoGVHD

Animal procedures were approved by the “Services Vétérinaires de la Préfecture de Police de Paris” and by the “Comité d’Ethique en matière d’Expérimentation Animale Paris Descartes (CEEA 34)” under the number APAFIS#23742-2017091815321774 v9, Université Paris Descartes, Paris, France. All appropriate procedures were performed in the animal facility (registration number A75-15-34) and followed to ensure animal welfare. Eight- to 12-week-old male NSG mice (bred in house or purchased from Charles River) were intraperitoneally injected with 25 mg/kg busulfan (Merck) on days -2 and -1 before injection of 5 x 10^6^ HLA-A2+ PBMCs into the tail vein with or without 5 x 10^6^ CAR-Tregs injected retro-orbitally via the venous sinus. Saline-injected mice served as controls. CAR-Tregs were generated from three different healthy donors. GVHD was scored based on weight, hunching, fur properties, diarrhea and skin integrity, with 0 to 1 point per category as described (Naserian et al., 2018). On the indicated days and after isoflurane anesthesia supplemented with tetracaine analgesia, peripheral blood from the venous sinus was harvested and centrifuged, and the plasma was collected and frozen before cytokine measurement. Then, the erythrocytes were lysed (RBC lysis buffer, Ozyme), and the leukocytes were evaluated by FACS. When a mouse reached a score of 4, it was sacrificed by cervical dislocation, and the spleen was collected for FACS analysis, whereas the lungs and liver were harvested for histology. Mouse tissues were fixed in 4% paraformaldehyde and paraffin embedded. Liver and lung sections (4 µm) were stained with hematoxylin and eosin, scanned (Nanozoomer 2.0, Hamamatsu) and blindly read by two independent researchers with NDPview software (Hamamatsu).

### Luciferase assay

To evaluate CAR-Treg homing *in vivo*, sorted Tregs (CD4+CD45RO+CD25^hi^CD127lo) were activated as described above. Two days later, the cells were transduced with either a CD28 CAR lentivirus or a 4-1BB CAR lentivirus at a MOI of 30 together with luciferase-mCherry-lentivirus at an MOI of 410. The lentiviral plasmid encoding a firefly luciferase protein (pEFS.eFFLY.mCherry) was kindly provided by Dr. Matthias Titeux. After 7 days of culture, double-transduced mCherry+EGFR+ Tregs expressing the CAR (or MOCK, the negative control) and luciferase were sorted before restimulation as described in Figure 1C. On day 18 of culture, 5 x 10^6^ luciferase-Tregs and 5 x 10^6^ human allogeneic HLA-A2+ PBMCs were injected intravenously into conditioned NSG mice. For bioluminescence imaging, D-luciferin potassium salt (150 mg/kg, Perkin Elmer) was injected intraperitoneally before anesthesia with isoflurane, and images were acquired within 20 minutes on an IVIS Spectrum CT (Perkin Elmer). Data were analyzed with Living Image software (IVIS Imaging Systems), and the BLI and X-ray superimposition signals were quantified after 3D bioluminescence imaging tomography (DLIT). Using coregistration with the Automatic Mouse Atlas, a region of interest (ROI) corresponding to the spleen was determined, and the absolute number of cells was obtained using *in vitro* calibration with corresponding cells.

### xCELLigence cytotoxicity assay

To evaluate the cytotoxic potential of CAR-Tregs, the viability of HLA-A2+ and HLA-class I-deficient endothelial cell lines was monitored every 15 minutes for 10 hours by electrical impedance measurement with an xCELLigence RTCA MP instrument (ACEA Biosciences). In each E-plate (ACEA Biosciences) well, 1 x 10^4^ HLA-A2+ or HLA-class I-deficient endothelial cells were seeded. After 15 hours, 2 x 10^4^ CAR-Tregs or CD8+ CAR-T cells were added to the culture. As a cytotoxicity control, CD8+ T cells were transduced 2 days after activation with the CAR construct at an MOI of 40 and incubated for 18 hours for transduction. On day 5 posttransduction, CD8+ EGFRt+ cells were sorted using a FACSAria II. CD8 CAR-T cells were cultured in complete medium supplemented with IL-2 (100 UI/ml).

The cell indices (CIs) were normalized to the reference value (measured just prior to adding CAR-Tregs to the culture). The normalized cell index in experimental wells was normalized over that of the control wells containing the endothelial cell lines only.

### Specific CAR-mediated activation in Jurkat RT3-T3.5 cells

Jurkat J.RT3-T3.5 cells (ATCC; TIB-153) transduced with either a CD28 CAR vector or a 4-1BB CAR vector were cocultured overnight with irradiated (35 Gy) HLA-A2/A28-positive or HLA-A2/A28-negative human splenocytes at a cell ratio of 1:1. After 20 hours of incubation, the cells were stained with an HLA-A2 MHC pentamer and anti-human EGFR antibody to monitor CAR expression; 7-aminoactinomycin D (7AAD) and an anti-human CD69 antibody were all used. Activation was defined as the upregulation of CD69 expression. Cross-reactivity was assessed against 8 different donors and up to 30 different allogeneic class I-HLA molecules.

### Statistical analysis

Results are presented as the mean+/-SEM or median for continuous variables. Frequencies of categorical variables are presented as numbers and percentages. Analyses were performed with GraphPad Prism software (version 8.00; GraphPad Software). For statistical comparisons of *in vitro* data, we used the nonparametric two-tailed Mann-Whitney test for comparisons of two groups. For survival comparisons, the log-rank test was used. P values <0.05 were considered significant.

## ABBREVIATIONS

CAR: Chimeric Antigen Receptor
CD: Cluster of differentiation
CSD: Costimulatory domains
CTLA4: Cytotoxic T-lymphocyte antigen 4
EC: Endothelial Cells
EFs: Elongation Factor 1-α shortened
EGFRt: truncated Epidermal Growth Factor Receptor
FOXP3: Forkhead-box protein 3
FVD: Fixable Viability Dye
GMP: Good Manufacturing Practice
GVHD: Graft-versus-host disease
MFI: Mean Fluorescence Intensity
MOI: Multiplicity of Infection
mTOR: mechanistic Target Of Rapamycin
mTORC: mTOR complex
PBMCs: Peripheral Blood Mononuclear Cells
PGE2: Prostaglandin E2
qPCR: quantitative Polymerase Chain Reaction
RFI: Ratio of fluorescence intensity
scFv: Single-chain variable fragment
T2A: Thosea asigna virus 2A
Tconvs: Conventional T cells
TNFRSF: TNF Receptor Superfamily
Tregs: T regulatory cells
TSDR: Treg-Specific Demethylated Region
VCN: Vector Copy Number

---

**Figure S1:**
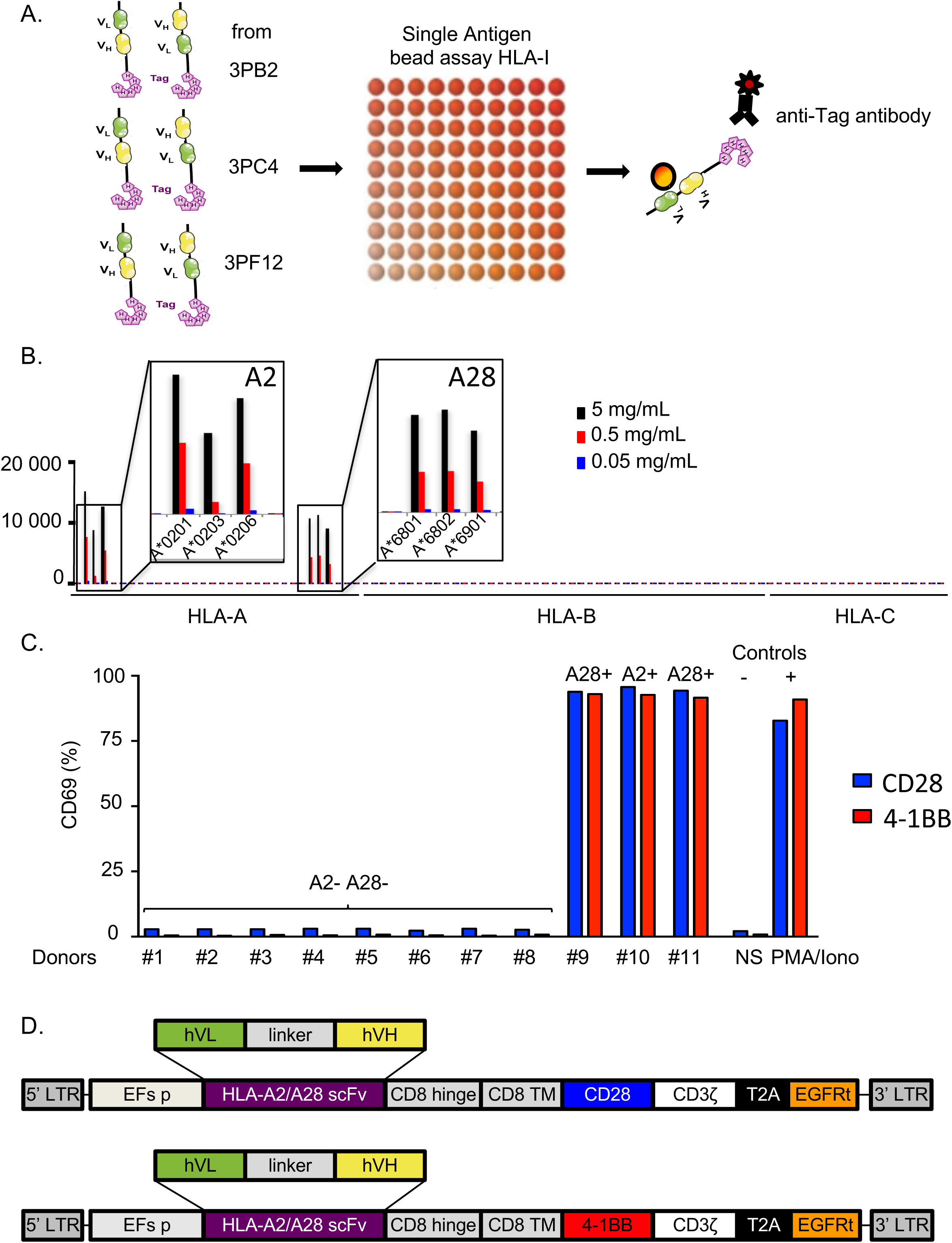
Selection of specific scFvs for CD28 and 4-1BB CAR constructs. A-B. Six tagged scFvs were tested in the Luminex single-antigen bead assay (class I HLA) at different concentrations, and specificity was assessed after PE-conjugated anti-Tag staining. A. Schematic design of the experiment. B. Histogram representing the specificity of selected scFvs at different concentrations. C. Histogram representing the frequency of activated CAR-J.RT3-T3.5 cells (CD69-positive cells) after 24 h of coculture with splenocytes according to HLA type. D. CD28 and 4-1BB CAR constructs with the same framework and an scFv against HLA-A2 and HLA-A28.

**Figure S2:**
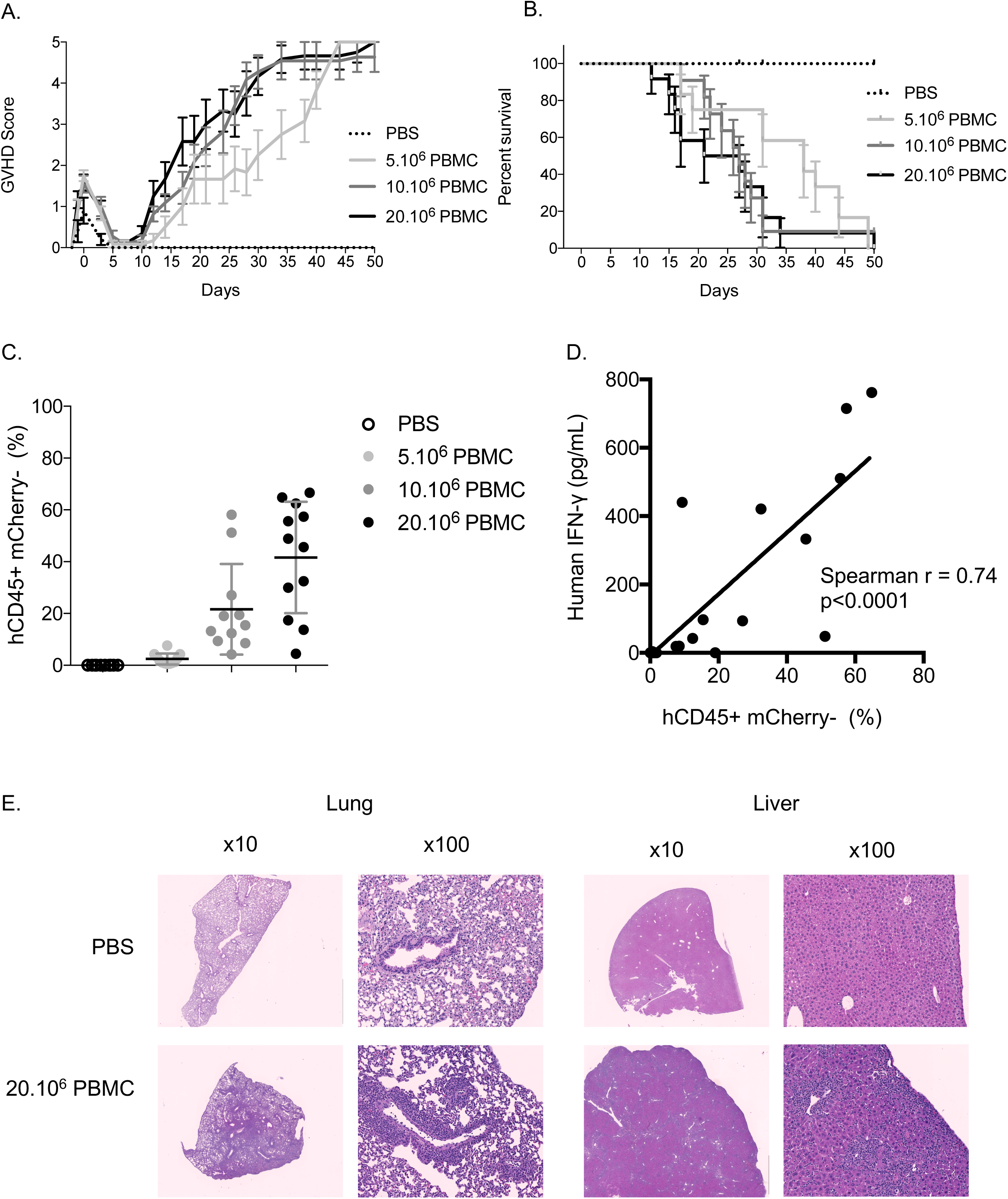
XenoGVHD model using a busulfan conditioning regimen. Eight- to 12-week-old male NSG mice were conditioned with busulfan on day -2 and day -1 before IV injection with the indicated number of HLA-A2+ PBMCs. Mice were weighed and scored for GVHD thrice weekly and bled weekly for flow cytometry analysis. A. GVHD score. B. Survival curves. C. At 10 days post-cell injection, the proportion of hCD45+ cells within the hCD45+ and mCD45+ cell populations was determined by flow cytometry. D. At 10, 17 and 24 days post-cell injection, the concentration of human IFN gamma in the plasma was measured by a cytometry bead array and correlated with human chimerism. E. Representative anatomopathological analysis of lesions in the liver and lungs performed on day 30 in NSG mice injected with 20 x 10^6^ HLA-A2+ PBMCs. The mean+/-SD is represented. N=3 mice per group from at least three independent experiments and three different donors.

**Figure S3:**
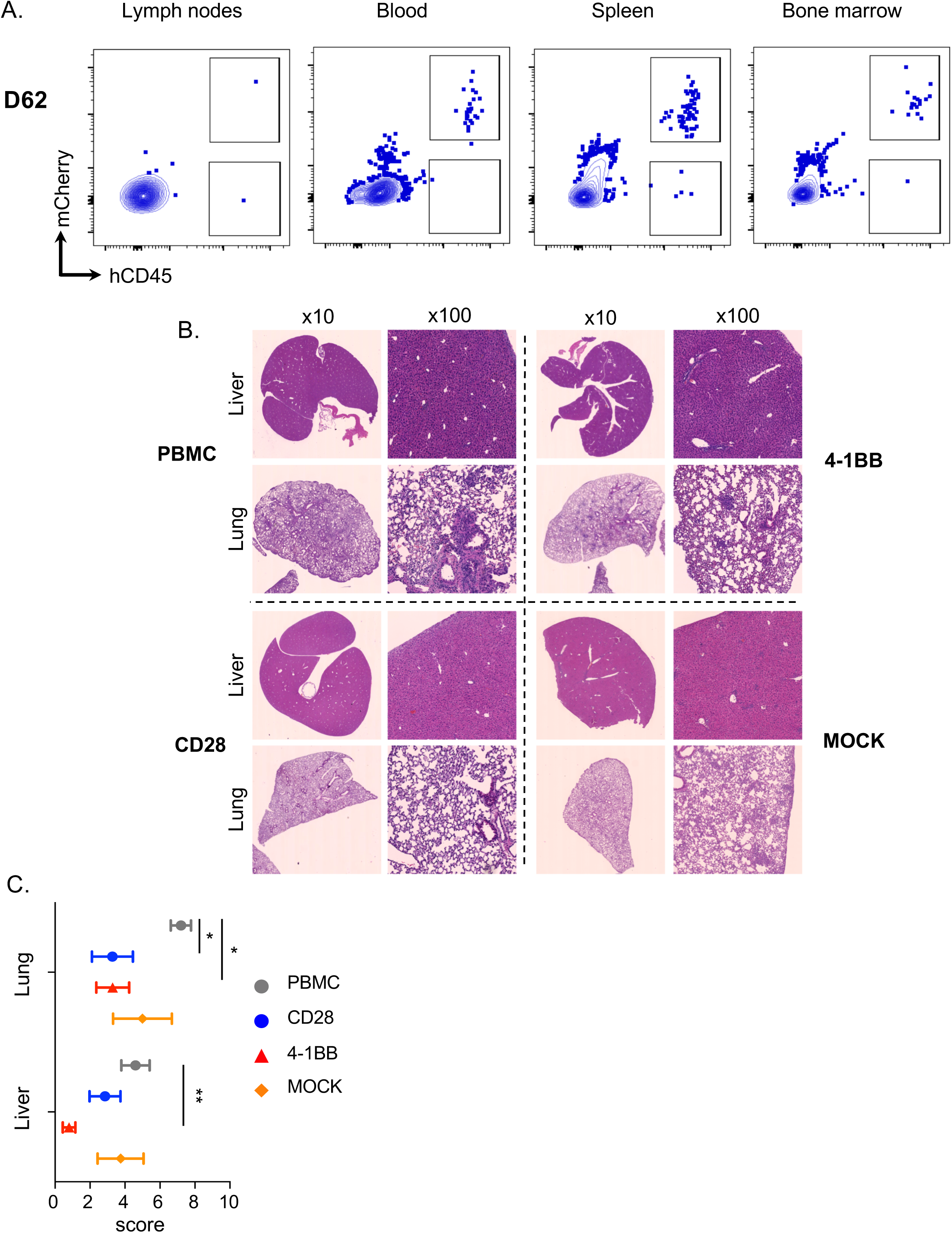
CAR-Treg detection in the xenoGVHD model. A. Representative contour plots of murine and human cells in the blood, spleen and bone marrow at sacrifice (days 60-62). B-C. Histologic analysis of lesions in the liver and lungs at sacrifice (either when mice reached a GVHD score>4 or on day 60 corresponding, to the end of experiment) in NSG mice injected with 5×10^6^ HLA-A2+ PBMCs followed by retro-orbital injection of 5×10^6^ of the indicated type of Tregs. C. Representative pathological features of infiltrates in the liver and lungs. C. Histological scores. The mean+/-SEM represents at least two independent experiments and two different donors. Mann-Whitney tests. *p<0.05, **p<0.01.

## ACKNOWLEDGMENTS

We thank S. Berissi and colleagues at the Plateforme d’histologie et morphologie du petit animal; O. Pellé and colleagues at the Plateforme de cytométrie; E. Panafieu and colleagues at the Laboratoire d’Expérimentation Animale et Transgénèse of the Structure Fédérative de Recherche Necker, Paris; and Gisèle Froment, Didier Nègre and Caroline Costa at the lentivector production facility/SFR BioSciences Gerland, Lyon Sud (UMS3444/US8). BL was supported by the IMAGINE Institute and the DIM-Thérapie Génique/Région Ile-de-France. AM was supported by the Assistance-Publique Hôpitaux de Paris. SC and TB were supported by the Fondation Emmanuel Boussard. The study was also funded by the Fondation Recherche Medicale, Fondation Centaure, Fondation Day-Solvay, Sandoz Pharmaceuticals, Agence de la Biomédecine and Agence Nationale de la Recherche. The IVIS spectrum used in this study was purchased with a grant from the DIM-Thérapie Génique/Région Ile-de-France.

## AUTHOR CONTRIBUTIONS

B.L., A.M., S.C., T.B. and J.Z. conceived and designed the experiments and wrote the paper. B.L., A.M., S.C. and T.B. performed the experiments and analyzed the data. K.V. and B.S. performed the TSDR analysis and analyzed the data. M.T. kindly provided the mCherry-Luciferase plasmid and helped with IVIS Spectrum bioluminescence analysis. T.B., M.D. and H.V. blindly read histological scores. E.S. provided help with the CAR constructs. N.P. provided reagents and insights for immunometabolism analysis. E.M. performed immunoblot experiments, and E.M. and S.L. analyzed the data. J-L.T. performed the Luminex analysis. D.A., C. L, E.M., S.L., M. C, and I.A. contributed to the discussion and manuscript editing.

## REFERENCES

Ajina, A., and J. Maher. 2018. Strategies to Address Chimeric Antigen Receptor Tonic Signaling. Mol. Cancer Ther. 17:1795–1815. doi:10.1158/1535-7163.MCT-17-1097.

Amir, E.D., K.L. Davis, M.D. Tadmor, E.F. Simonds, J.H. Levine, S.C. Bendall, D.K. Shenfeld, S. Krishnaswamy, G.P. Nolan, and D. Pe’er. 2013. viSNE enables visualization of high dimensional single-cell data and reveals phenotypic heterogeneity of leukemia. Nat. Biotechnol. 31:545–552. doi:10.1038/nbt.2594.

Arvey, A., J. van der Veeken, G. Plitas, S.S. Rich, P. Concannon, and A.Y. Rudensky. 2015. Genetic and epigenetic variation in the lineage specification of regulatory T cells. Elife. 4:e07571. doi:10.7554/eLife.07571.

Bacher, P., F. Heinrich, U. Stervbo, M. Nienen, M. Vahldieck, C. Iwert, K. Vogt, J. Kollet, N. Babel, B. Sawitzki, C. Schwarz, S. Bereswill, M.M. Heimesaat, G. Heine, G. Gadermaier, C. Asam, M. Assenmacher, O. Kniemeyer, A.A. Brakhage, F. Ferreira, M. Wallner, M. Worm, and A. Scheffold. 2016. Regulatory T Cell Specificity Directs Tolerance versus Allergy against Aeroantigens in Humans. Cell. 167:1067–1078.e16. doi:10.1016/j.cell.2016.09.050.

Battaglia, M., A. Stabilini, B. Migliavacca, J. Horejs-Hoeck, T. Kaupper, and M.-G. Roncarolo. 2006. Rapamycin promotes expansion of functional CD4+CD25+FOXP3+ regulatory T cells of both healthy subjects and type 1 diabetic patients. J. Immunol. 177:8338–47. doi:10.4049/jimmunol.177.12.8338.

Blank, C.U., W.N. Haining, W. Held, P.G. Hogan, A. Kallies, E. Lugli, R.C. Lynn, M. Philip, A. Rao, N.P. Restifo, A. Schietinger, T.N. Schumacher, P.L. Schwartzberg, A.H. Sharpe, D.E. Speiser, E.J. Wherry, B.A. Youngblood, and D. Zehn. 2019. Defining “T cell exhaustion”. Nat. Rev. Immunol. 19:665–674. doi:10.1038/s41577-019-0221-9.

Boardman, D.A., C. Philippeos, G.O. Fruhwirth, M.A.A. Ibrahim, R.F. Hannen, D. Cooper, F.M. Marelli-Berg, F.M. Watt, R.I. Lechler, J. Maher, L.A. Smyth, and G. Lombardi. 2017. Expression of a Chimeric Antigen Receptor Specific for Donor HLA Class I Enhances the Potency of Human Regulatory T Cells in Preventing Human Skin Transplant Rejection. Am. J. Transplant. 17:931– 943. doi:10.1111/ajt.14185.

Boroughs, A.C., R.C. Larson, B.D. Choi, A.A. Bouffard, L.S. Riley, E. Schiferle, A.S. Kulkarni, C.L. Cetrulo, D. Ting, B.R. Blazar, S. Demehri, and M. V Maus. 2019. Chimeric antigen receptor costimulation domains modulate human regulatory T cell function. JCI insight. 5. doi:10.1172/jci.insight.126194.

Cazaux, M., C.L. Grandjean, F. Lemaître, Z. Garcia, R.J. Beck, I. Milo, J. Postat, J.B. Beltman, E.J. Cheadle, and P. Bousso. 2019. Single-cell imaging of CAR T cell activity in vivo reveals extensive functional and anatomical heterogeneity. J. Exp. Med. 216:1038–1049. doi:10.1084/jem.20182375.

Chi, H. 2012. Regulation and function of mTOR signalling in T cell fate decisions. Nat. Rev. Immunol. 12:325–338. doi:10.1038/nri3198.

Cosentino, M., A.M. Fietta, M. Ferrari, E. Rasini, R. Bombelli, E. Carcano, F. Saporiti, F. Meloni, F. Marino, and S. Lecchini. 2007. Human CD4+CD25+ regulatory T cells selectively express tyrosine hydroxylase and contain endogenous catecholamines subserving an autocrine/paracrine inhibitory functional loop. Blood. 109:632–642. doi:10.1182/blood-2006-01-028423.

Dawson, N.A., C. Lamarche, R.E. Hoeppli, P. Bergqvist, V.C. Fung, E. McIver, Q. Huang, J. Gillies, M. Speck, P.C. Orban, J.W. Bush, M. Mojibian, and M.K. Levings. 2019. Systematic testing and specificity mapping of alloantigen-specific chimeric antigen receptors in regulatory T cells. JCI insight. 4. doi:10.1172/jci.insight.123672.

Dawson, N.A.J., I. Rosado-Sánchez, G.E. Novakovsky, V.C.W. Fung, Q. Huang, E. McIver, G. Sun, J. Gillies, M. Speck, P.C. Orban, M. Mojibian, and M.K. Levings. 2020. Functional effects of chimeric antigen receptor co-receptor signaling domains in human regulatory T cells. Sci. Transl. Med. 12. doi:10.1126/scitranslmed.aaz3866.

Bin Dhuban, K., E. d’Hennezel, E. Nashi, A. Bar-Or, S. Rieder, E.M. Shevach, S. Nagata, and C.A. Piccirillo. 2015. Coexpression of TIGIT and FCRL3 identifies Helios+ human memory regulatory T cells. J. Immunol. 194:3687–3696. doi:10.4049/jimmunol.1401803.

DuPage, M., G. Chopra, J. Quiros, W.L. Rosenthal, M.M. Morar, D. Holohan, R. Zhang, L. Turka, A. Marson, and J.A. Bluestone. 2015. The Chromatin-Modifying Enzyme Ezh2 Is Critical for the Maintenance of Regulatory T Cell Identity after Activation. Immunity. 42:227–238. doi:10.1016/j.immuni.2015.01.007.

Fernández-Ramos, A.A., C. Marchetti-Laurent, V. Poindessous, S. Antonio, C. Petitgas, I. Ceballos-Picot, P. Laurent-Puig, S. Bortoli, M.-A. Loriot, and N. Pallet. 2017. A comprehensive characterization of the impact of mycophenolic acid on the metabolism of Jurkat T cells. Sci. Rep. 7:10550. doi:10.1038/s41598-017-10338-6.

Ferraro, A., A.M. D’Alise, T. Raj, N. Asinovski, R. Phillips, A. Ergun, J.M. Replogle, A. Bernier, L. Laffel, B.E. Stranger, P.L. De Jager, D. Mathis, and C. Benoist. 2014. Interindividual variation in human T regulatory cells. Proc. Natl. Acad. Sci. U. S. A. 111:E1111–20. doi:10.1073/pnas.1401343111.

Frigault, M.J., J. Lee, M.C. Basil, C. Carpenito, S. Motohashi, J. Scholler, O.U. Kawalekar, S. Guedan, S.E. McGettigan, A.D. Posey, S. Ang, L.J.N. Cooper, J.M. Platt, F.B. Johnson, C.M. Paulos, Y. Zhao, M. Kalos, M.C. Milone, C.H. June, and C.H. June. 2015. Identification of chimeric antigen receptors that mediate constitutive or inducible proliferation of T cells. Cancer Immunol. Res. 3:356–67. doi:10.1158/2326-6066.CIR-14-0186.

Galgani, M., V. De Rosa, A. La Cava, and G. Matarese. 2016. Role of Metabolism in the Immunobiology of Regulatory T Cells. J. Immunol. 197:2567–2575. doi:10.4049/jimmunol.1600242.

Gerriets, V.A., R.J. Kishton, M.O. Johnson, S. Cohen, P.J. Siska, A.G. Nichols, M.O. Warmoes, A.A. De Cubas, N.J. MacIver, J.W. Locasale, L.A. Turka, A.D. Wells, and J.C. Rathmell. 2016. Foxp3 and Toll-like receptor signaling balance T reg cell anabolic metabolism for suppression. Nat. Immunol. 17:1459–1466. doi:10.1038/ni.3577.

Gomes-Silva, D., M. Mukherjee, M. Srinivasan, G. Krenciute, O. Dakhova, Y. Zheng, J.M.S. Cabral, C.M. Rooney, J.S. Orange, M.K. Brenner, and M. Mamonkin. 2017. Tonic 4-1BB Costimulation in Chimeric Antigen Receptors Impedes T Cell Survival and Is Vector-Dependent. Cell Rep. 21:17– 26. doi:10.1016/j.celrep.2017.09.015.

Hauck, F., C. Randriamampita, E. Martin, S. Gerart, N. Lambert, A. Lim, J. Soulier, Z. Maciorowski, F. Touzot, D. Moshous, P. Quartier, S. Heritier, S. Blanche, F. Rieux-Laucat, N. Brousse, I. Callebaut, A. Veillette, C. Hivroz, A. Fischer, S. Latour, and C. Picard. 2012. Primary T-cell immunodeficiency with immunodysregulation caused by autosomal recessive LCK deficiency. J. Allergy Clin. Immunol. 130:1144–1152.e11. doi:https://doi.org/10.1016/j.jaci.2012.07.029.

He, T., S. Liu, S. Chen, J. Ye, X. Wu, Z. Bian, and X. Chen. 2018. The p38 MAPK Inhibitor SB203580 Abrogates Tumor Necrosis Factor-Induced Proliferative Expansion of Mouse CD4(+)Foxp3(+) Regulatory T Cells. Front. Immunol. 9:1556. doi:10.3389/fimmu.2018.01556.

Hoffmann, P., R. Eder, T.J. Boeld, K. Doser, B. Piseshka, R. Andreesen, and M. Edinger. 2006. Only the CD45RA+ subpopulation of CD4+CD25 high T cells gives rise to homogeneous regulatory T-cell lines upon in vitro expansion. Blood. 108:4260–4267. doi:10.1182/blood-2006-06-027409.

Huynh, A., M. Dupage, B. Priyadharshini, P.T. Sage, J. Quiros, C.M. Borges, N. Townamchai, V.A. Gerriets, J.C. Rathmell, A.H. Sharpe, J.A. Bluestone, and L.A. Turka. 2015. Control of PI(3) kinase in Tregcells maintains homeostasis and lineage stability. Nat. Immunol. 16:188–196. doi:10.1038/ni.3077.

Iwano, S., M. Sugiyama, H. Hama, A. Watakabe, N. Hasegawa, T. Kuchimaru, K.Z. Tanaka, M. Takahashi, Y. Ishida, J. Hata, S. Shimozono, K. Namiki, T. Fukano, M. Kiyama, H. Okano, S. Kizaka-Kondoh, T.J. McHugh, T. Yamamori, H. Hioki, S. Maki, and A. Miyawaki. 2018. Single-cell bioluminescence imaging of deep tissue in freely moving animals. Science. 359:935–939. doi:10.1126/science.aaq1067.

Kawalekar, O.U., R.S. O’Connor, J.A. Fraietta, L. Guo, S.E. McGettigan, A.D. Posey, P.R. Patel, S. Guedan, J. Scholler, B. Keith, N.W. Snyder, I.A. Blair, M.C. Milone, C.H. June, M.C. Milone, and C.H. June. 2016. Distinct Signaling of Coreceptors Regulates Specific Metabolism Pathways and Impacts Memory Development in CAR T Cells. Immunity. 44:380–390. doi:10.1016/j.immuni.2016.01.021.

Kendal, A.R., Y. Chen, F.S. Regateiro, J. Ma, E. Adams, S.P. Cobbold, S. Hori, and H. Waldmann. 2011. Sustained suppression by Foxp3+ regulatory T cells is vital for infectious transplantation tolerance. J. Exp. Med. 208:2043–53. doi:10.1084/jem.20110767.

Kim, H.-J., R.A. Barnitz, T. Kreslavsky, F.D. Brown, H. Moffett, M.E. Lemieux, Y. Kaygusuz, T. Meissner, T.A.W. Holderried, S. Chan, P. Kastner, W.N. Haining, and H. Cantor. 2015. Stable inhibitory activity of regulatory T cells requires the transcription factor Helios. Science. 350:334–339. doi:10.1126/science.aad0616.

Latour, S., G. Gish, C.D. Helgason, R.K. Humphries, T. Pawson, and A. Veillette. 2001. Regulation of SLAM-mediated signal transduction by SAP, the X-linked lymphoproliferative gene product. Nat. Immunol. 2:681–690. doi:10.1038/90615.

Livak, K.J., and T.D. Schmittgen. 2001. Analysis of Relative Gene Expression Data Using Real-Time Quantitative PCR and the 2−ΔΔCT Method. Methods. 25:402–408. doi:https://doi.org/10.1006/meth.2001.1262.

Long, A.H., W.M. Haso, J.F. Shern, K.M. Wanhainen, M. Murgai, M. Ingaramo, J.P. Smith, A.J. Walker, M.E. Kohler, V.R. Venkateshwara, R.N. Kaplan, G.H. Patterson, T.J. Fry, R.J. Orentas, and C.L. Mackall. 2015. 4-1BB costimulation ameliorates T cell exhaustion induced by tonic signaling of chimeric antigen receptors. Nat. Med. 21:581–590. doi:10.1038/nm.3838.

Lowther, D.E., B.A. Goods, L.E. Lucca, B.A. Lerner, K. Raddassi, D. van Dijk, A.L. Hernandez, X. Duan, M. Gunel, V. Coric, S. Krishnaswamy, J.C. Love, and D.A. Hafler. 2016. PD-1 marks dysfunctional regulatory T cells in malignant gliomas. JCI Insight. 1. doi:10.1172/jci.insight.85935.

Lu, L., J. Barbi, and F. Pan. 2017. The regulation of immune tolerance by FOXP3. Nat. Rev. Immunol. 17:703–717. doi:10.1038/nri.2017.75.

MacDonald, K.G., R.E. Hoeppli, Q. Huang, J. Gillies, D.S. Luciani, P.C. Orban, R. Broady, and M.K. Levings. 2016. Alloantigen-specific regulatory T cells generated with a chimeric antigen receptor. J. Clin. Invest. 126:1413–24. doi:10.1172/JCI82771.

Menk, A. V, N.E. Scharping, D.B. Rivadeneira, M.J. Calderon, M.J. Watson, D. Dunstane, S.C. Watkins, and G.M. Delgoffe. 2018. 4-1BB costimulation induces T cell mitochondrial function and biogenesis enabling cancer immunotherapeutic responses. J. Exp. Med. 215:1091–1100. doi:10.1084/jem.20171068.

Milone, M.C., J.D. Fish, C. Carpenito, R.G. Carroll, G.K. Binder, D. Teachey, M. Samanta, M. Lakhal, B. Gloss, G. Danet-Desnoyers, D. Campana, J.L. Riley, S.A. Grupp, and C.H. June. 2009. Chimeric receptors containing CD137 signal transduction domains mediate enhanced survival of T cells and increased antileukemic efficacy in vivo. Mol. Ther. 17:1453–64. doi:10.1038/mt.2009.83.

Miyara, M., Y. Yoshioka, A. Kitoh, T. Shima, K. Wing, A. Niwa, C. Parizot, C. Taflin, T. Heike, D. Valeyre, A. Mathian, T. Nakahata, T. Yamaguchi, T. Nomura, M. Ono, Z. Amoura, G. Gorochov, and S. Sakaguchi. 2009. Functional Delineation and Differentiation Dynamics of Human CD4+ T Cells Expressing the FoxP3 Transcription Factor. Immunity. 30:899–911. doi:10.1016/j.immuni.2009.03.019.

Naserian, S., M. Leclerc, A. Thiolat, C. Pilon, C. Le Bret, Y. Belkacemi, S. Maury, F. Charlotte, and J.L. Cohen. 2018. Simple, Reproducible, and Efficient Clinical Grading System for Murine Models of Acute Graft-versus-Host Disease. Front. Immunol. 9:10. doi:10.3389/fimmu.2018.00010.

Newton, R., B. Priyadharshini, and L.A. Turka. 2016. Immunometabolism of regulatory T cells. Nat. Immunol. 17:618–625. doi:10.1038/ni.3466.

Nowak, A., D. Lock, P. Bacher, T. Hohnstein, K. Vogt, J. Gottfreund, P. Giehr, J.K. Polansky, B. Sawitzki, A. Kaiser, J. Walter, and A. Scheffold. 2018. CD137+CD154− Expression As a Regulatory T Cell (Treg)-Specific Activation Signature for Identification and Sorting of Stable Human Tregs from In Vitro Expansion Cultures. Front. Immunol. 9:199. doi:10.3389/fimmu.2018.00199.

Noyan, F., K. Zimmermann, M. Hardtke-Wolenski, A. Knoefel, E. Schulde, R. Geffers, M. Hust, J. Huehn, M. Galla, M. Morgan, A. Jokuszies, M.P. Manns, and E. Jaeckel. 2017. Prevention of Allograft Rejection by Use of Regulatory T Cells With an MHC-Specific Chimeric Antigen Receptor. Am. J. Transplant. 17:917–930. doi:10.1111/ajt.14175.

Pierini, A., B.P. Iliopoulou, H. Peiris, M. Pérez-Cruz, J. Baker, K. Hsu, X. Gu, P.-P. Zheng, T. Erkers, S.-W. Tang, W. Strober, M. Alvarez, A. Ring, A. Velardi, R.S. Negrin, S.K. Kim, and E.H. Meyer. 2017. T cells expressing chimeric antigen receptor promote immune tolerance. JCI Insight. 2. doi:10.1172/jci.insight.92865.

Procaccini, C., F. Carbone, D. Di Silvestre, F. Brambilla, V. De Rosa, M. Galgani, D. Faicchia, G. Marone, D. Tramontano, M. Corona, C. Alviggi, A. Porcellini, A. La Cava, P. Mauri, and G. Matarese. 2016. The Proteomic Landscape of Human Ex Vivo Regulatory and Conventional T Cells Reveals Specific Metabolic Requirements. Immunity. 44:406–421. doi:10.1016/j.immuni.2016.01.028.

Sagoo, P., N. Ali, G. Garg, F.O. Nestle, R.I. Lechler, and G. Lombardi. 2011. Human regulatory T cells with alloantigen specificity are more potent inhibitors of alloimmune skin graft damage than polyclonal regulatory T cells. Sci. Transl. Med. 3:83ra42. doi:10.1126/scitranslmed.3002076.

Salomon, B., D.J. Lenschow, L. Rhee, N. Ashourian, B. Singh, A. Sharpe, and J.A. Bluestone. 2000. B7/CD28 costimulation is essential for the homeostasis of the CD4+CD25+ immunoregulatory T cells that control autoimmune diabetes. Immunity. 12:431–40. doi:10.1016/s1074-7613(00)80195-8.

Sasidharan Nair, V., M.H. Song, and K.I. Oh. 2016. Vitamin C Facilitates Demethylation of the Foxp3 Enhancer in a Tet-Dependent Manner. J. Immunol. 196:2119–31. doi:10.4049/jimmunol.1502352.

Savage, T.M., B.A. Shonts, A. Obradovic, S. Dewolf, S. Lau, J. Zuber, M.T. Simpson, E. Berglund, J. Fu, S. Yang, S.-H. Ho, Q. Tang, L.A. Turka, Y. Shen, and M. Sykes. 2018. Early expansion of donor-specific Tregs in tolerant kidney transplant recipients. JCI Insight. 3. doi:10.1172/jci.insight.124086.

Sawitzki, B., P.N. Harden, P. Reinke, A. Moreau, J.A. Hutchinson, D.S. Game, Q. Tang, E.C. Guinan, M. Battaglia, W.J. Burlingham, I.S.D. Roberts, M. Streitz, R. Josien, C.A. Böger, C. Scottà, J.F. Markmann, J.L. Hester, K. Juerchott, C. Braudeau, B. James, L. Contreras-Ruiz, J.B. van der Net, T. Bergler, R. Caldara, W. Petchey, M. Edinger, N. Dupas, M. Kapinsky, I. Mutzbauer, N.M. Otto, R. Öllinger, M.P. Hernandez-Fuentes, F. Issa, N. Ahrens, C. Meyenberg, S. Karitzky, U. Kunzendorf, S.J. Knechtle, J. Grinyó, P.J. Morris, L. Brent, A. Bushell, L.A. Turka, J.A. Bluestone, R.I. Lechler, H.J. Schlitt, M.C. Cuturi, S. Schlickeiser, P.J. Friend, T. Miloud, A. Scheffold, A. Secchi, K. Crisalli, S.-M. Kang, R. Hilton, B. Banas, G. Blancho, H.-D. Volk, G. Lombardi, K.J. Wood, and E.K. Geissler. 2020. Regulatory cell therapy in kidney transplantation (The ONE Study): a harmonised design and analysis of seven non-randomised, single-arm, phase 1/2A trials. Lancet (London, England). 395:1627–1639. doi:10.1016/S0140-6736(20)30167-7.

Schoenbrunn, A., M. Frentsch, S. Kohler, J. Keye, H. Dooms, B. Moewes, J. Dong, C. Loddenkemper, J. Sieper, P. Wu, C. Romagnani, N. Matzmohr, and A. Thiel. 2012. A converse 4-1BB and CD40 ligand expression pattern delineates activated regulatory T cells (Treg) and conventional T cells enabling direct isolation of alloantigen-reactive natural Foxp3+ Treg. J. Immunol. 189:5985–94. doi:10.4049/jimmunol.1201090.

Sugimoto, N., and Y.-J. Liu. 2013. DUSP4 Stabilizes FOXP3 Expression In Human Regulatory T Cells. Blood. 122:3473. doi:10.1182/blood.V122.21.3473.3473.

Vasanthakumar, A., Y. Liao, P. Teh, M.F. Pascutti, A.E. Oja, A.L. Garnham, R. Gloury, J.C. Tempany, T. Sidwell, E. Cuadrado, P. Tuijnenburg, T.W. Kuijpers, N. Lalaoui, L.A. Mielke, V.L. Bryant, P.D. Hodgkin, J. Silke, G.K. Smyth, M.A. Nolte, W. Shi, and A. Kallies. 2017. The TNF Receptor Superfamily-NF-κB Axis Is Critical to Maintain Effector Regulatory T Cells in Lymphoid and Non-lymphoid Tissues. Cell Rep. 20:2906–2920. doi:10.1016/j.celrep.2017.08.068.

Vignali, D.A.A., L.W. Collison, and C.J. Workman. 2008. How regulatory T cells work. Nat. Rev. Immunol. 8:523–32. doi:10.1038/nri2343.

Watanabe, N., Y.-H. Wang, H.K. Lee, T. Ito, Y.-H. Wang, W. Cao, and Y.-J. Liu. 2005. Hassall’s corpuscles instruct dendritic cells to induce CD4+CD25+ regulatory T cells in human thymus. Nature. 436:1181–1185. doi:10.1038/nature03886.

Watkins, N.A., C. Brown, C. Hurd, C. Navarrete, and W.H. Ouwehand. 2000. The isolation and characterisation of human monoclonal HLA-A2 antibodies from an immune V gene phage display library. Tissue Antigens. 55:219–228. doi:10.1034/j.1399-0039.2000.550305.x.

Wieczorek, G., A. Asemissen, F. Model, I. Turbachova, S. Floess, V. Liebenberg, U. Baron, D. Stauch, K. Kotsch, J. Pratschke, A. Hamann, C. Loddenkemper, H. Stein, H.D. Volk, U. Hoffmuller, A. Grutzkau, A. Mustea, J. Huehn, C. Scheibenbogen, and S. Olek. 2009. Quantitative DNA Methylation Analysis of FOXP3 as a New Method for Counting Regulatory T Cells in Peripheral Blood and Solid Tissue. Cancer Res. 69:599–608. doi:10.1158/0008-5472.CAN-08-2361.

Xu, T., K.M. Stewart, X. Wang, K. Liu, M. Xie, J. Kyu Ryu, K. Li, T. Ma, H. Wang, L. Ni, S. Zhu, N. Cao, D. Zhu, Y. Zhang, K. Akassoglou, C. Dong, E.M. Driggers, and S. Ding. 2017. Metabolic control of TH17 and induced Treg cell balance by an epigenetic mechanism. Nature. 548:228– 233. doi:10.1038/nature23475.

Yue, X., S. Trifari, T. Äijö, A. Tsagaratou, W.A. Pastor, J.A. Zepeda-Martínez, C.-W.J. Lio, X. Li, Y. Huang, P. Vijayanand, H. Lähdesmäki, and A. Rao. 2016. Control of Foxp3 stability through modulation of TET activity. J. Exp. Med. 213:377–397. doi:10.1084/jem.20151438.

Zeng, H., K. Yang, C. Cloer, G. Neale, P. Vogel, and H. Chi. 2013. mTORC1 couples immune signals and metabolic programming to establish T(reg)-cell function. Nature. 499:485–490. doi:10.1038/nature12297.

Zhang, R., A. Huynh, G. Whitcher, J. Chang, J.S. Maltzman, and L.A. Turka. 2013. An obligate cell-intrinsic function for CD28 in Tregs. J. Clin. Invest. 123:580–93. doi:10.1172/JCI65013.

Zhao, Z., M. Condomines, S.J.C. van der Stegen, F. Perna, C.C. Kloss, G. Gunset, J. Plotkin, and M. Sadelain. 2015. Structural Design of Engineered Costimulation Determines Tumor Rejection Kinetics and Persistence of CAR T Cells. Cancer Cell. 28:415–428. doi:10.1016/j.ccell.2015.09.004.

Zuber, J., M. Viguier, F. Lemaitre, V. Senée, N. Patey, G. Elain, F. Geissmann, F. Fakhouri, L. Ferradini, C. Julier, and A. Bandeira. 2007. Severe FOXP3+ and naïve T lymphopenia in a non-IPEX form of autoimmune enteropathy combined with an immunodeficiency. Gastroenterology. 132:1694–704. doi:10.1053/j.gastro.2007.02.034.

